# Chromatinization Modulates Topoisomerase II Processivity

**DOI:** 10.1101/2023.10.03.560726

**Authors:** Jaeyoon Lee, Meiling Wu, James T. Inman, Gundeep Singh, Seong ha Park, Joyce H. Lee, Robert M. Fulbright, Yifeng Hong, Joshua Jeong, James M. Berger, Michelle D. Wang

## Abstract

Type IIA topoisomerases are essential DNA processing enzymes that must robustly and reliably relax DNA torsional stress *in vivo*. While cellular processes constantly create different degrees of torsional stress, how this stress feeds back to control type IIA topoisomerase function remains obscure. Using a suite of single-molecule approaches, we examined the torsional impact on supercoiling relaxation of both naked DNA and chromatin by eukaryotic topoisomerase II (topo II). We observed that topo II was at least ∼ 50-fold more processive on plectonemic DNA than previously estimated, capable of relaxing > 6000 turns. We further discovered that topo II could relax supercoiled DNA prior to plectoneme formation, but with a ∼100-fold reduction in processivity; strikingly, the relaxation rate in this regime decreased with diminishing torsion in a manner consistent with the capture of transient DNA loops by topo II. Chromatinization preserved the high processivity of the enzyme under high torsional stress. Interestingly, topo II was still highly processive (∼ 1000 turns) even under low torsional stress, consistent with the predisposition of chromatin to readily form DNA crossings. This work establishes that chromatin is a major stimulant of topo II function, capable of enhancing function even under low torsional stress.

## INTRODUCTION

Topoisomerases are ubiquitous enzymes required for solving a variety of topological problems resulting from the double-helical structure of DNA^1, 2, 3, 4^. Type IIA topoisomerases are of particular interest due to their ability to both relax supercoiled DNA and decatenate DNA molecules^5, 6^. These enzymes act on DNA through a strand-passage mechanism, in which they hydrolyze ATP to pass one segment of DNA (the transfer or T-segment) through a transient, enzyme-mediated double-stranded break in another segment (the gate or G-segment)^5, 6^. Notably, this action requires the enzyme to capture both the G- and T-segments, necessitating that two DNA segments be in close proximity and form a crossing prior to strand passage.

Eukaryotic type IIA topoisomerases, also known as topoisomerase II (topo II), are vital for the resolution of torsional stress during transcription and replication^2, 3, 7^. *In vivo*, torsional stress is dynamic, varying over both space and time. During transcription, RNA polymerase progression generates DNA supercoiling, which increases as the transcription level increases^8^. Torsion accumulated near active genes can impact cellular functions thousands of base pairs away due to supercoil diffusion^9^. Similarly, during DNA replication, the degree of supercoiling near the replication fork varies over time, with torsional stress increasing towards termination^10^. Thus, for topo II to properly support supercoiling homeostasis and cell viability^11, 12, 13, 14^, it must be able to relieve dynamically varying levels of torsional stress, yet whether or how the enzyme adjusts its activity in response to varying levels of torsional stress is not fully understood. Previous biochemical and single-molecule studies have successfully elucidated many aspects of topo II activity on naked, plectonemically supercoiled (or ‘buckled’) DNA under high torsional stress^6, 15, 16, 17, 18, 19, 20^. However, torsion can also accumulate in “pre-buckled” DNA prior to buckling, and there has yet to be any clear assessment of topo II action on such a substrate. Even less is known about the impact of chromatinization on enzyme processivity and supercoil relaxation rate, limiting our understanding of the enzyme in its *in vivo* context.

Questions relating to the torsional response of topo II have been challenging to address due to constraints in experimental approaches. Previous techniques have been limited to studies of the topoisomerase-mediated relaxation of DNA that are undergoing time-varying changes in topological state. In particular, single-molecule studies of topo II reported to date have only measured the relaxation of naked buckled DNA^15, 18, 19^. In addition, while type IIA topoisomerases from various organisms have been shown to preferentially bind to supercoiled DNA over relaxed DNA^21, 22, 23^, how the processivity and strand passage rate of topo II depends on torsional stress has remained elusive. Chromatin presents a particularly significant set of challenges for studying topo activity and, to date, direct measurements of topo II’s biophysical properties on chromatin have not been possible.

In the present work, we developed single-molecule assays that enable the direct measurement of topo II activity on varying degrees of torsionally strained DNA in both buckled (plectonemically supercoiled) and pre-buckled (non-plectonemically supercoiled) regimes. We then applied these assays to chromatin to characterize the impact of chromatinization and torsion on topo II activity. Taken together, our results show that chromatinization modulates topo II activity by enhancing processivity, enabling the enzyme to efficiently resolve torsional stress even prior to DNA buckling.

## RESULTS

### Quantitative definition of torsional stress

To understand how topo II responds to different degrees of torsional stress, it is important to first quantitatively define this metric. Torque in DNA resists rotation/twist, so torsion can be characterized by torque (analogous to linear stress being characterized by force, which resists translocation). Using an angular optical trap (AOT) setup (Fig. 1a), we have directly measured the torque necessary to add turns to DNA as a function of the number of turns added (DNA supercoiling state) using methods we established previously^24, 25, 26, 27, 28^. As shown in Fig. 1b, when turns are added to DNA held under a given force, torque (torsion) in DNA increases until the DNA buckles to extrude a plectoneme. Before the buckling transition, a significant fraction of the added turns is converted to DNA twist, while after the buckling transition, torque no longer increases with the further addition of turns, and all additional turns are converted to DNA writhe (Fig. 1b, c). Thus, torsional stress increases with added turns prior to buckling but remains constant after buckling.

**Figure 1.**
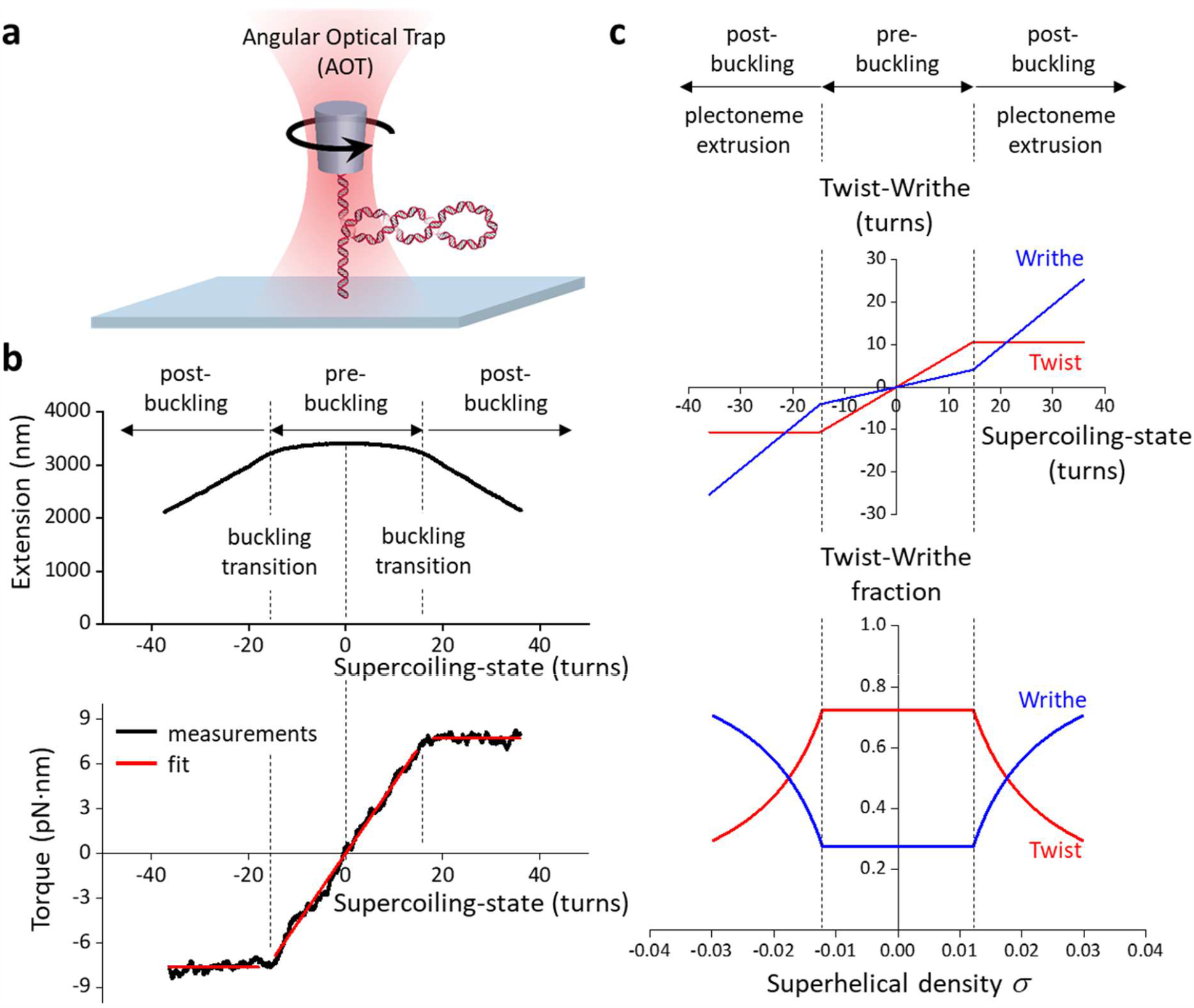
Torsional response of naked DNA measured by the angular optical trap (AOT). (**a**) Experimental configuration for measuring DNA torsional response on the AOT. A 12.7 kb DNA molecule containing a random sequence was torsionally constrained between a birefringent quartz cylinder and the coverslip surface of a microscope sample chamber. The AOT simultaneously measured and controlled the DNA extension, force, rotation, and torque. (**b**) Extension and torque versus turns added to DNA under constant force. Data were averaged from *N* = 22 DNA molecules held at 0.5 pN. In the pre-buckled regime, torque increased linearly as turns were added. The linear fit within the pre-buckled regime (middle red line) yields an effective twist persistence length of 78.9 ± 0.3 nm, consistent with previous measurements within measurement uncertainty^28^. At the buckling transitions (vertical dashed black lines), DNA underwent phase transitions to form plectonemes. The torque plateaued with further plectoneme extrusion at +7.7 ± 0.2 pN·nm (mean ± s.d) for the (+) supercoiled DNA and -7.6 ± 0.3 pN·nm for the (-) supercoiled DNA (horizontal red lines). (**c**) Partitioning of twist and writhe. The DNA supercoiling state is defined by the turns added to DNA, or the change in the linking number ∆*Lk* = *Lk* − *Lk*_0_, with *Lk*_0_ being the linking number of torsionally relaxed DNA. ∆*Lk* is partitioned into a change in twist ∆*Tw* (torsion) and a change in writhe ∆*Wr*: ∆*Lk* = ∆*Tw* + ∆*Wr*. The superhelical density *σ* is used to characterize the degree of supercoiling: 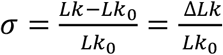. Using the fit parameters from (**b**), the partitioning of added turns into the twist and writhe was obtained. In the pre-buckling regime, the added turns partition mostly into twist, which increases linearly with turns; the partitioning of twist was calculated from the ratio of the effective twist persistence length, 78.9 nm, and the intrinsic twist persistence length, 109 nm^28^. Beyond the buckling transition, all added turns partition into writhe, thus maintaining a constant torque.

### Topo II is highly processive on buckled DNA

We first systematically investigated topo II processivity on buckled and pre-buckled DNA (Fig. 2). To enable direct measurement of topo II processivity and relaxation rate on a buckled DNA in a constant topological state, we developed a single-molecule constant-extension method to clamp the DNA extension (i.e., maintain a constant superhelical density) as topo II continuously relaxed the substrate. We implemented this method on a custom-built magnetic tweezers (MT) instrument (Fig. 2a; Supplementary Fig. 1; Methods). To measure the activity of a single topo II enzyme on a DNA molecule, we pre-incubated the sample chamber with a low concentration of yeast topo II (0.5 pM) and flushed out excess topo from the chamber (Methods). We also conducted control experiments to show that our measurements should be predominately from a single topo II (Methods). After topo II binding, we turned on the extension clamp and kept the DNA in a post-buckled state via magnet rotation, whose rate was equal and opposite to the topo II relaxation rate to counteract the activity of the enzyme and thus served as an excellent readout of topo II activity. Notably, this constant-extension method effectively creates an infinitely renewable substrate maintained in a constant buckled state throughout topo II action, permitting the measurement of topo processivity well beyond that attainable in a conventional supercoiling relaxation experiment while also guaranteeing that the DNA always remains well within the buckled regime.

**Figure 2.**
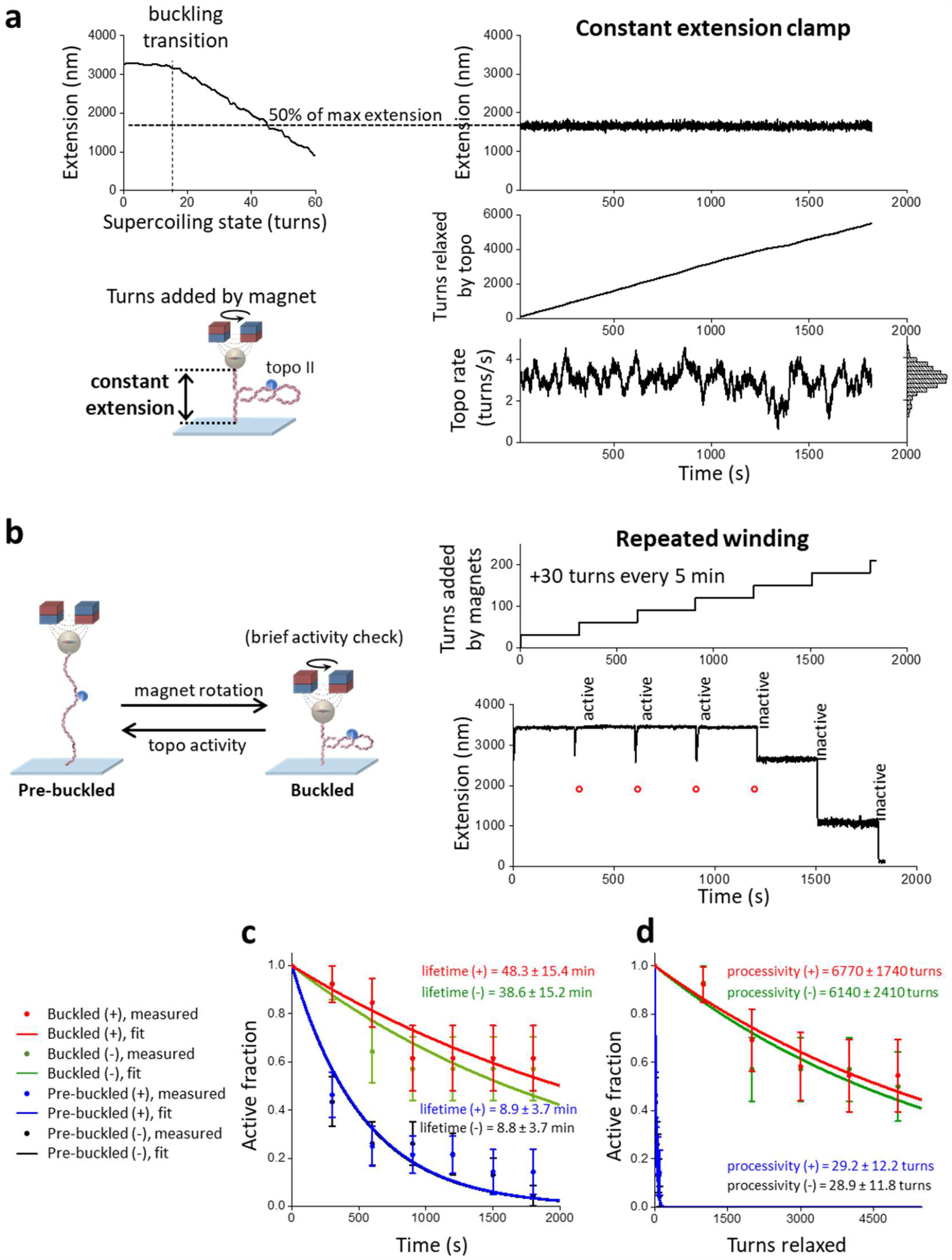
Plectonemes stabilize topo II activity on DNA. (**a**) Measurements of topo II activity lifetime, processivity, and relaxation rate on a buckled DNA in a constant topological state. 12.7 kb DNA molecules containing a random sequence were torsionally constrained between a magnetic bead and the coverslip surface of a microscope sample chamber, and initial extension-supercoiling state relations were measured under 0.5 pN (top left). A DNA molecule showing topo II activity was selected, wound to 50% of its zero-turn extension into a buckled state, then held at this extension for 30 min via magnet rotation counteracting topo activity (right). The topo relaxation rate was calculated from the magnet turns filtered by a 20 s window; the rate distribution is shown as a histogram (bottom right). (**b**) Measurements of topo II activity lifetime and processivity on pre-buckled DNA. +30 turns were added every 5 min to DNA molecules under 0.5 pN by rapid rotation of the magnets (40 turns/s) (top right). After each winding step, DNA molecules with active topo II were identified by increases in extension; molecules without topo II activity remained at a constant extension until the next winding step. A representative trace with topo II activity for the first four winding steps is shown (bottom right). Red circles indicate expected extension if topo II was not able to relax pre-buckled DNA; if the extension immediately after a winding step (tip of a downward spike) was above a red circle, then topo II had relaxed DNA into the pre-buckled regime during the previous relaxation step. (**c**) Activity lifetime of topo II on (+) buckled (red, *N* = 13), (-) buckled (green, *N* = 14), (+) pre-buckled (blue, *N* = 28), and (-) pre-buckled DNA (black, *N* = 23); error bars represent counting errors. These data were fit to exponential curves to estimate the lifetimes. (**d**) Processivity of topo II on (+) and (-) buckled DNA and (+) and (-) pre-buckled DNA from the same set of traces considered in Fig. 2c; error bars represent counting errors. These data were fit to exponential curves to estimate the processivities.

Unexpectedly, we discovered that budding yeast topo II was extremely processive on buckled DNA. Fig. 2a shows a typical trace of a constant-extension measurement with a single yeast topo II molecule that remained active for 30 min, at which point the experiment was manually terminated. During this time, the topo II molecule removed 5500 turns from the DNA. Strikingly, this processivity is about 50 times greater than those determined from previous measurements for human topo IIα^18^. To determine whether this difference might be due to the innate properties of distinct species of enzyme, we also applied the constant-extension method to human topo IIα and human IIβ (Fig. 3). As with yeast topo II, both human topo II enzymes were also found to be highly processive, capable of relaxing thousands of turns without dissociation. Thus, high processivity on buckled DNA appears to be a relatively general feature of eukaryotic type IIA topoisomerases.

**Figure 3.**
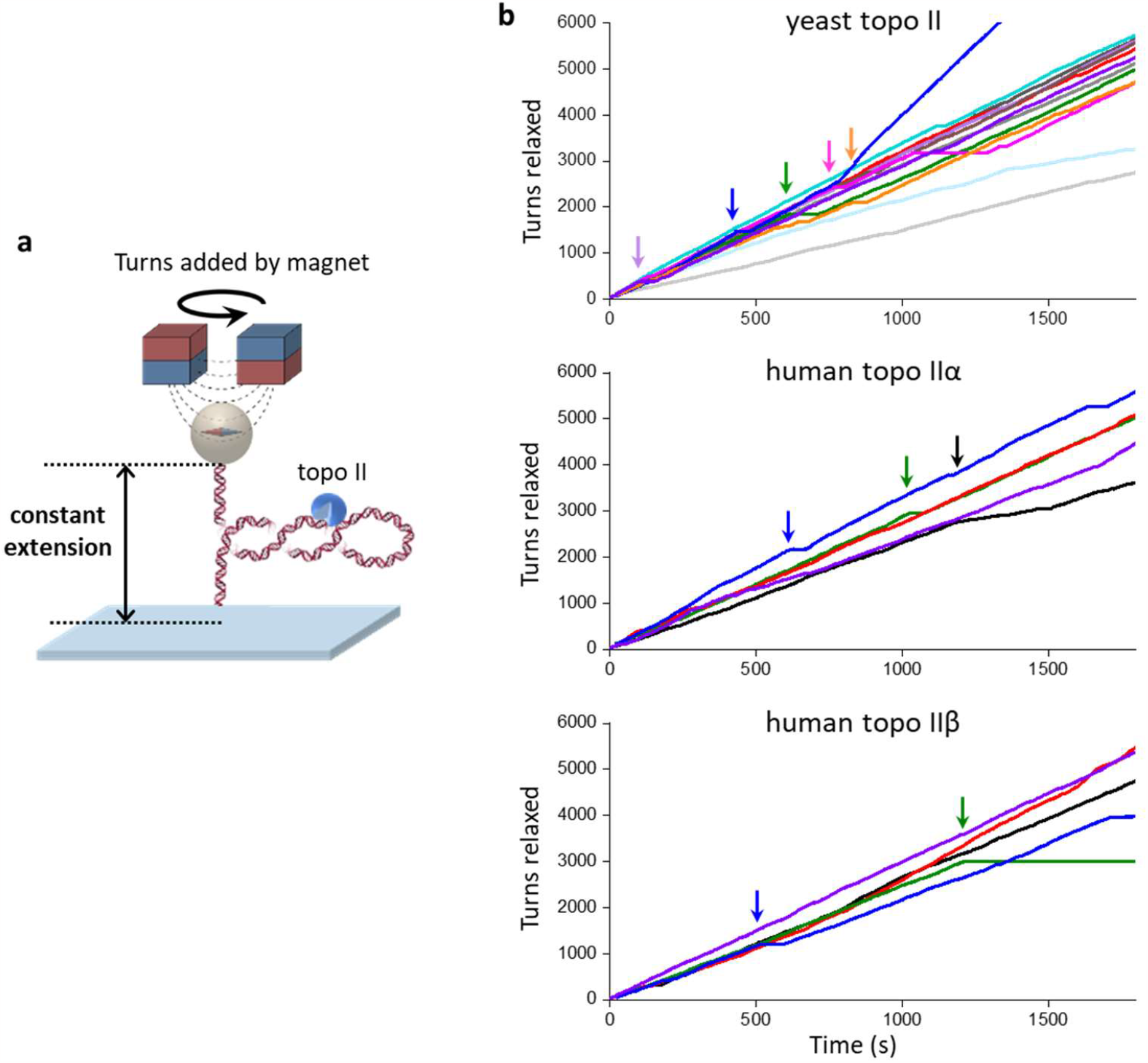
Human topo IIα and IIβ are also highly processive on buckled DNA. (**a**) The constant-extension experimental configuration, as shown in Fig. 2a (reshown here for clarity). All experiments shown in this figure were conducted on the 12.7 kb DNA clamped at 50% extension under (+) supercoiling. (**b**) Topo II activity traces of turns relaxed vs. time: yeast topo II (top panel, *N* = 13, same traces as those used in Figs. 2c and 2d), human topo IIα (middle panel, *N* = 5), and human topo IIβ (bottom panel, *N* = 5). Each color represents a measurement from a single DNA molecule, with an arrow of the same color indicating the first occurrence when the topo II paused for > 30 s. For the dark blue trace of yeast topo II, topo II relaxation paused with duration > 30 s at 430 s (and thus classified as no longer active at that time) and later showed a doubling of the rate beginning at 770 s, likely due to tandem relaxation by two topo II molecules.

In addition to the processivity, each trace provided a measurement of the instantaneous relaxation rate from a single topo II molecule. The rate of relaxation in this trace was homogeneous at 3.0 turns/s on average, albeit with some modest variation (coefficient of variation of 0.19) that could reflect the stochastic nature of topo II activity (Fig. 2a, bottom right). This variation is also consistent with enzyme-to-enzyme variations (Supplementary Fig. 1e). Because topo II changes the linking number (Lk) of DNA in steps of 2 with each reaction cycle, this value indicates that the enzyme is catalyzing 1-2 rounds of strand passage per second, a number in good accord with previous biochemical measurements^29^.

### Topo II is less processive on pre-buckled DNA

Although topo II activity is extremely stable and processive on buckled DNA, it has been unclear whether topo II is similarly active on pre-buckled DNA, which is under lower torsional stress and does not contain a plectoneme to provide suitable G- and T-segments in close proximity for topo II to act on. Unfortunately, detecting the relaxation of pre-buckled DNA is challenging, as any strand passage event by topo II will lead to only a minimal DNA extension change in the pre-buckled regime. To circumvent this limitation, we performed a “repeated winding” experiment to measure topo II activity on pre-buckled DNA (Fig. 2b). For this approach, we repeatedly checked for topo II activity by periodically adding a small number of turns to the DNA with magnet rotation to bring the molecules slightly into the buckled regime. If topo II was active, the extension of the tether increased as the topo II relaxed the newly added supercoils. Thus, the subsequent rewinding step enabled us to check if topo II was active during the preceding pre-buckled DNA state.

The repeated winding study revealed that topo II is clearly capable of relaxing pre-buckled DNA. Fig. 2b shows a typical trace from this experiment, in which +30 additional turns were added every 5 minutes by rapidly rotating the magnets at 40 turns/s. Topo II remained active for three additional rewinding steps after the initial winding step, as demonstrated by the increasing extension after those rewinding steps. The extension after each of those rewinding steps corresponded to the expected extension if, at the start of the rewinding step, the DNA were near the fully relaxed state instead of at the buckling transition. This result indicates that in the time between successive rewinding steps, topo II relaxed all plectonemic supercoils and then continued to relax the supercoiling in the pre-buckled DNA to a near-fully relaxed state. To our knowledge, processive relaxation of pre-buckled DNA by a type IIA topoisomerase to near completion has not been previously reported. It is worth nothing that the DNA was buckled for only a small duration (0.75 s) during the rewinding step in which topo II relaxed at most 2 turns, so relaxation of the added turns occurred primarily in the pre-buckled state. In addition, the 5 min of topo II relaxation in between rewinding steps was not enough for complete relaxation of the DNA (Supplementary Fig. 2), so topo II was continuously relaxing supercoiling as long as it remained active. Thus, the activity lifetime measured by this experiment should accurately reflect that of topo II on pre-buckled DNA, i.e., for the trace shown in Fig. 2b, topo II retained activity on the pre-buckled DNA for 15-20 min.

We next compared statistics from multiple traces obtained with (+) supercoiled buckled and pre-buckled DNA to examine the effect of torsional stress on topo II processivity (Fig. 2c, d). On buckled DNA, we did not observe any permanent loss of activity before 30 min, although 40% of traces contained long pauses (> 30 s duration) that may be due to the bound topo II becoming temporarily inactive before being active again (Fig.3). It is also possible that the bound topo II became inactive or dissociated and relaxation resumed after the binding of another topo II molecule, but we expect such events to be extremely rare (Methods). Nonetheless, when calculating the fraction of traces with continued topo activity (“active fraction”) as a function of time, we conservatively classified individual traces as being “active” only until the first such pause to avoid overestimating the processivity. Fitting the active fraction over time to an exponential yielded a lower bound on the mean activity lifetime of 48.3 ± 15.4 min. We also obtained a lower bound on the processivity of topo II in turns by considering the active fraction versus turns relaxed, which yielded 6770 ± 1740 turns, or 3390 ± 870 enzymatic turnovers (1 enzymatic turnover = 2 turns relaxed). This lower bound on the processivity (> 6000 turns) far exceeds values suggested by other single-molecule methods for similar enzymes^15, 18, 19^. Our results with human topo IIα and IIβ showing high processivity (Fig. 3) suggest that this discrepancy arises from differences in methodology, with the constant-extension method ensuring that topo II acts only on buckled DNA so that its extreme processivity on this substrate is observable.

In contrast with the relaxation of buckled DNA, our repeated winding experiment showed that topo II associates much less stably with pre-buckled DNA, with a mean activity lifetime of 8.9 ± 3.7 min. This result may be a slight overestimate because topo II that was initially bound to DNA could have dissociated and been replaced by another topo II in between successive rewinding steps, which would be indistinguishable from the same topo remaining bound in our experiment. However, we again expect such events to be rare because free topo II was flushed out of our sample chamber (Methods), and this possibly slightly overestimated value is still far smaller than the activity lifetime of topo II on buckled DNA. Since the DNA extension immediately after each rewinding step indicated the DNA supercoiling state in the pre-buckled regime before rewinding (Supplementary Fig. 2), we could also estimate the processivity of topo II in the pre-buckled regime by plotting the active fraction as a function of the cumulative number of turns applied to the DNA in the pre-buckled regime. Fitting to an exponential yielded a mean processivity of 29.2 ± 12.2 turns, a number far smaller than the lower bound on the processivity of topo II on buckled DNA (6770 ± 1740 turns). To assess whether the activity lifetime or processivity of topo II show any dependency on the chirality of the torsional stress (i.e., over-vs. under-winding), we then repeated the same activity lifetime and processivity measurements by applying (-) supercoiling DNA. Our results indicate that there is no significant dependence of topo II activity lifetimes and processivities on the chirality of supercoiling applied on naked DNA (Fig. 2c, d), in good agreement with a previous biochemical study^17^.

Taken together, our findings establish that topo II is much more stably associated with buckled DNA compared to on pre-buckled DNA. Topo II can relax many thousands of turns in a buckled DNA before losing activity, about 50 times greater than previous estimates. In contrast, on pre-buckled DNA, processivity is reduced by at least ∼100 fold. These findings may provide an explanation for previous studies which showed that type IIA topoisomerases preferentially interact with supercoiled DNA compared to relaxed DNA^21, 22, 23^ and indicate that the increased affinity could stem from a marked reduction in the dissociation rate.

### Topo II rate slows down on pre-buckled DNA

Torsional stress in a DNA substrate could influence not only topo II processivity but also the enzyme’s relaxation rate. We found that the relaxation rate of topo II was constant on a buckled DNA, independent of the plectonemic length (i.e., superhelical density) (Supplementary Fig. 3). On the other hand, while Fig. 2b shows that topo II can relax pre-buckled DNA, it was unclear how topo II relaxation rate might depend on the DNA superhelical density and torsion in this regime. We therefore developed a method to investigate the ability of topo II to relax DNA in the pre-buckled regime (Fig. 4a). For this method, we first buckled a DNA molecule by the addition of a fixed number of turns and monitored the DNA extension as it increased over time due to topo II supercoiling relaxation. Once the DNA extension reached the buckling transition, we allowed for a specified duration of further topo II relaxation into the pre-buckled regime. We then performed a “buckling check” by applying an additional +35 rewinding turns to bring the DNA back into the buckled state, whose extension provided the final supercoiling state of the preceding topo II relaxation step (Methods). We repeated this process of allowing topo II to relax pre-buckled DNA for different durations, varying from 2 s to 15 s, each followed by a DNA buckling check. Through this approach, we were able to track the relaxation of pre-buckled DNA over time. For each trace, this sequence was repeated either until topo II activity was lost, or up to a maximum of 5 times. It is worth noting that the rewinding was brief (0.875 s) with topo II relaxing ∼ 2.5 turns on average during each rewinding step (Methods).

**Figure 4.**
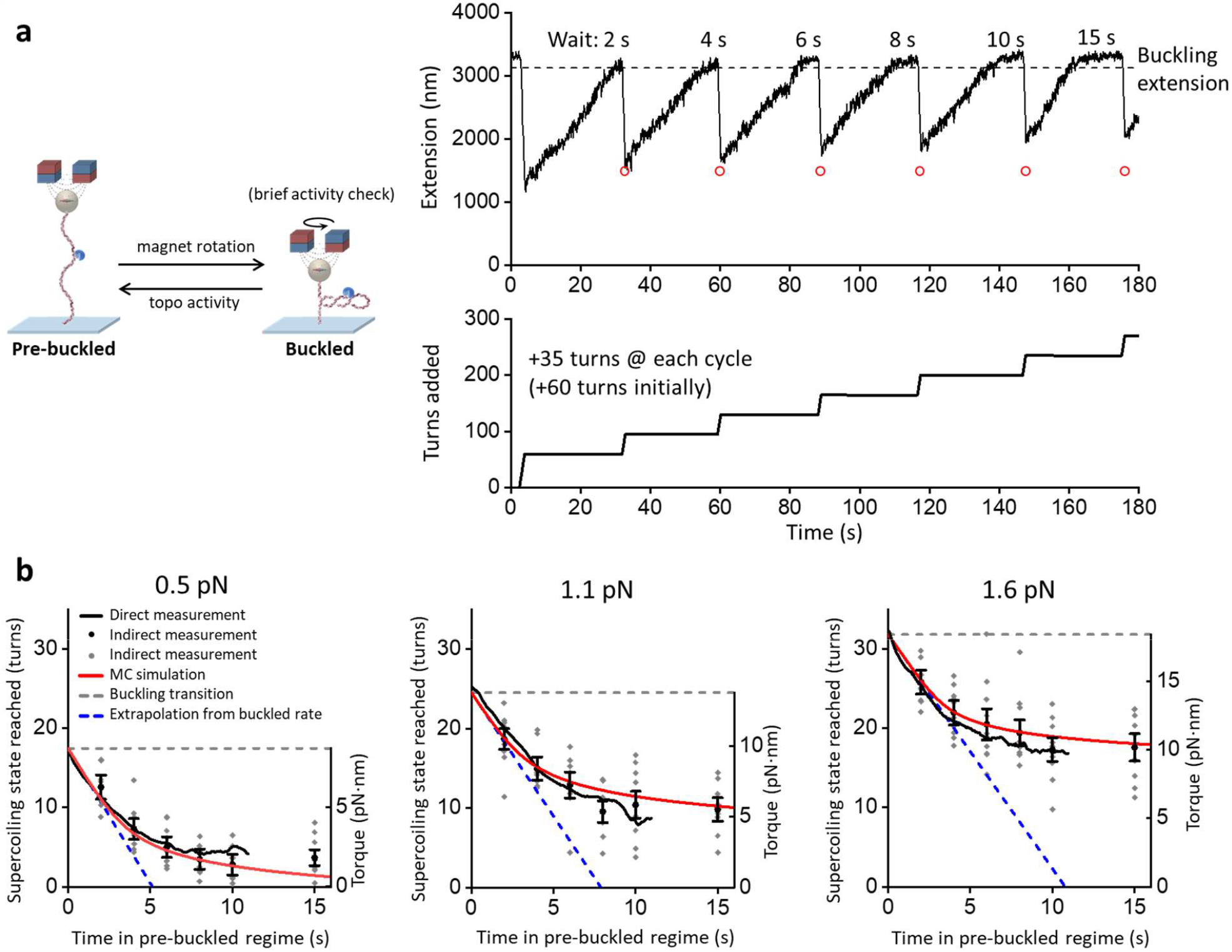
Topo II captures spontaneous loops in pre-buckled DNA. (**a**) Measurements of topo II relaxation of pre-buckled DNA. Initially, +60 turns were added by rapid rotation of the magnets (40 turns/s) to a 12.7 kb DNA molecule containing a random sequence and with an active topo II bound. Once the extension reached the buckling transition point due to topo II activity, a set duration of activity in the pre-buckled regime was allowed before +35 additional turns were added by the magnets. If the extension immediately after a winding step (tip of a downward spike) was above the red circle (expected extension after winding if relaxed only to the buckling transition), then topo II had relaxed DNA into the pre-buckled regime during the previous relaxation step. The process was repeated for different durations. (**b**) Supercoiling state reached by topo II relaxation into the pre-buckled regime versus time of topo II relaxing the pre-buckled regime under 0.5 pN (*N* = 10 traces), 1.1 pN (*N* = 9 traces), and 1.6 pN (*N* = 11 traces). We calculated the DNA supercoiling state using two methods. In the direct-measurement method, the relaxation time courses during the 15 s wait shown in (a) was used to directly determine supercoiling state versus wait time in the pre-buckled state (Methods). In the indirect-measurement method, the supercoiling state was determined based on the shift in the extension-supercoiling state relation before and after topo relaxation in the pre-buckled state for each trace (Methods). For the indirect method, we show a scatter plot of all data points (gray) along with their means and SEMs for each duration of topo II activity on pre-buckled DNA (black). The torque in DNA corresponding to a supercoiling state is shown as the right vertical axis based on previous torque measurements^28^ using methods similar to those of Fig. 1b. The dashed blue lines are linear extrapolations using the relaxation rates on buckled DNA (Methods) and project the supercoiling state progression if topo II had not slowed down in the pre-buckled regime. The red curves are theoretical results from a simple model assuming that topo II relaxation rate depends on the DNA loop formation rate, which we obtain from Monte-Carlo simulations, following a Michaelis-Menten type equation (Methods).

As shown in Fig. 4b, the relaxation of pre-buckled DNA by topo II was initially as efficient as the relaxation of buckled DNA. However, topo II activity slowed down sharply as the DNA torsion decreased before the DNA became completely relaxed. To check whether topo II activity stopped completely at the observed plateau, we performed a separate experiment that allowed topo II to relax pre-buckled DNA for a much longer time (10 min instead of 15 s) (Supplementary Fig. 2) and found that topo II relaxed the pre-buckled DNA even further but still not to completion, suggesting a slow rate towards full relaxation.

The above method provides an indirect measurement of the progression of topo II relaxation in the pre-buckled regime. To validate this method, we also directly converted the extension data to the supercoiling state of DNA versus time in the pre-buckled state using the DNA extension-supercoiling state relation (Methods). As shown in Fig. 4b, the supercoiling state determined using the direct method agrees with that from the indirect method within the uncertainty of the measurement. This agreement provides strong evidence that topo II relaxes into the pre-buckled state, although the relaxation rate slows down as the DNA torsion decreases.

### Topo II captures loops that form spontaneously in pre-buckled DNA

Based on our findings, we hypothesized that topo II relaxes pre-buckled DNA in the absence of a plectoneme by capturing spontaneous loops that form as the DNA configuration thermally fluctuates due to DNA bending and writhing^30, 31, 32, 33^. The incidence of such loops should depend on the DNA supercoiling state, with loops forming more often near or after the buckling transition than on more relaxed DNA^31^. As topo II relaxes DNA toward the fully relaxed state, spontaneous loops may become limiting, leading to the observed reduction in relaxation rate.

If this hypothesis is correct, then topo II should be less effective relaxing pre-buckled DNA that is held under a greater force, which is known to suppress spontaneous loop formation ^31, 32^. To test this prediction, we performed the same experiment under different forces (0.5 pN, 1.1 pN, and 1.6 pN) (Fig. 4b). We observed that the number of turns left in the DNA after 10-15 s of topo II activity in the pre-buckled regime increased with increasing force, consistent with a reduction in spontaneous loop formation at a higher force. Interestingly, at a given supercoiling state, topo II relaxation rate slightly decreased when DNA torsion (torque) increased with an increase in force from 0.5 pN to 1.6 pN, suggesting that changes in torsion alone are not sufficient to explain the variation in topo II relaxation rate.

Furthermore, spontaneous loop formation should decrease for a shorter DNA molecule, which disfavors thermally induced looping^31^. Thus, we repeated the experiment with DNA molecules that were half the length of our primary substrate and found that the relaxation of pre-buckled DNA is relatively slower on this shorter DNA (Supplementary Fig. 4). As a further check, we also considered a simple writhe model that has been previously used for topo II relaxation of post-buckled DNA by assuming that the topo II relaxation rate is only dependent on writhe according to the Michaelis-Menten relation^19^. We found that such a model cannot explain our measurements in the pre-buckled regime (Supplementary Fig. 5), suggesting that DNA writhing alone cannot explain topo II activity in this regime. Thus, while DNA writhe and the number of DNA loops are related, these two quantities are fundamentally not the same.

To relate topo II activity in a pre-buckled regime to DNA loop formation, we performed Monte Carlo simulations to compute the rate of DNA crossing formation as a function of DNA supercoiling state and force (Supplementary Fig. 6; Methods). For each supercoiling state under a given force, we first generated equilibrium configurations of pre-buckled DNA using the parameters shown in Supplementary Table 1. From these configurations, we then calculated the rate of DNA crossing formation and modeled the topo II relaxation rate by assuming it depends on the DNA crossing rate via a Michaelis-Menten type relationship (Methods). The results of these simulations show excellent agreement with our measurements at all forces (Fig. 4b). Because yeast topo II is known to introduce a bend in DNA^34, 35^, we also repeated the simulations by including a bend to examine how DNA bending may affect the DNA crossing rate. Simulations including such a bend also show good agreement with our measurements (Supplementary Fig. 7). Taken together, our data indicate that topo II relaxes pre-buckled DNA by capturing spontaneous DNA loops.

### Chromatinization enhances topo II processivity

Chromatin is the natural substrate of eukaryotic topo II *in vivo*. Although there is some evidence that the C-terminus of topo II can associate with histone proteins^36, 37^, the catalytic domains of topo II likely interact directly with the linker DNA that connects nucleosomes. Since chromatinization markedly alters the structural and torsional mechanical properties of DNA^27^, we speculate that the activity of topo II on chromatinized substrates may be significantly different from that on wholly naked DNA. In particular, chromatin exhibits an asymmetric extension-supercoiling state relation indicative of three distinct topological states^27^: two steep regimes on the left and right sides (which we consider to be the (-) and (+) buckling-like regimes, respectively), and a middle plateau regime with a slight negative slope (Fig. 5a, top). Each nucleosome contributes about +1.0 turns to the (+) width of the plateau, implying that each nucleosome can absorb about 1.0 turns of (+) supercoiling before buckling. The presence of the extension plateau indicates that chromatin can effectively buffer (+) torsional stress, absorbing significant (+) DNA supercoiling without torsional build-up^27, 38, 39^. Consistent with this, the measured torque under (+) supercoiling shows a greatly reduced torsional stiffness in the extension plateau regime (Fig. 5a, bottom).

**Figure 5.**
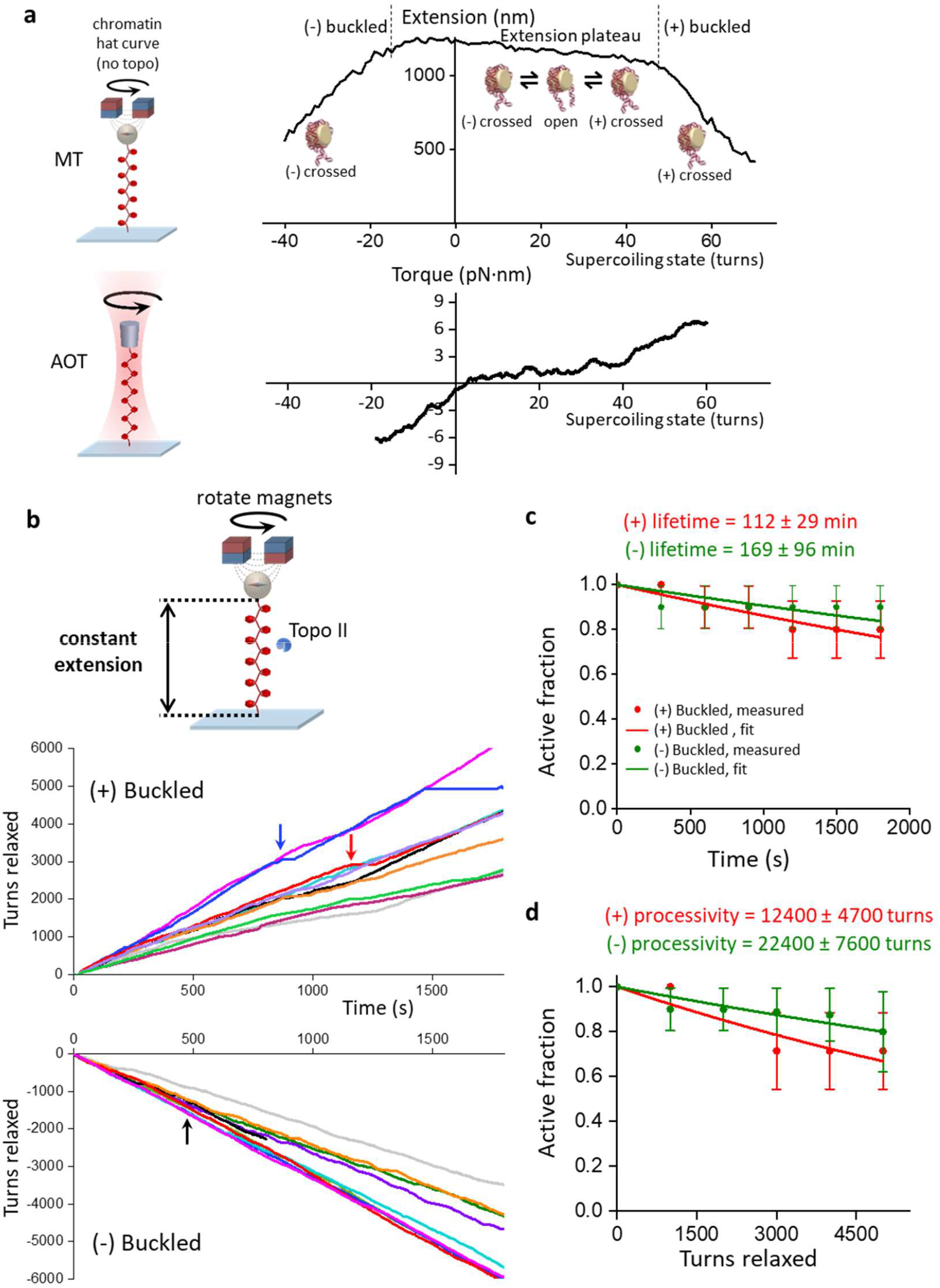
Chromatinization preserves high topo II processivity under high torsional stress. (**a**) Measurements of chromatin extension-supercoiling state relation before introducing topo II. Chromatin fibers were assembled on 12.7 kb DNA containing 64 repeats of a nucleosome-positioning element and were torsionally constrained between magnetic beads and the coverslip surface of a microscope sample chamber under 0.5 pN magnetic force. The top panel is a representative extension-supercoiling state relation for a chromatin fiber containing 48 nucleosomes. Shown in the bottom panel is the corresponding torque-supercoiling state relation previously measured using the angular optical trap^27^. (**b**) Topo II activity traces of turns relaxed vs. time. Data were acquired using the constant-extension method at 50% of the maximal extension of the chromatin fiber under 0.5 pN. These experiments were performed the same way as shown in Figs. 2a and 3 but with chromatin fibers instead of naked DNA. Each color represents a measurement from a single chromatin fiber, with an arrow of the same color indicating the first occurrence when the topo II paused for > 30 s. For the black trace under (-) buckling, topo II relaxation paused at 260 s, and the trace was manually terminated at 800 s when another free magnetic bead in the sample drifted over the tether and obstructed the bead position detection. (**c, d**) Activity lifetime and processivity of topo II on buckled chromatin under high (+) torsional stress (*N* = 10 traces) and high (-) torsional stress (*N* = 10 traces) from data shown in (**b**); error bars represent counting error. These data were fit with exponential curves to estimate the activity lifetimes and processivities.

To examine the effect of chromatinization on topo II activity under high torsional stress, we again used the constant-extension method (Figs. 2a and 3) but now with chromatin fibers containing ∼50 nucleosomes assembled on a DNA construct comprising 64 repeats of a nucleosome positioning element (Fig. 5b; Methods). For this measurement, we pre-bound single topo II enzymes to chromatin fibers that satisfied our selection criteria (Supplementary Fig. 8; Methods) and used the extension clamp to keep the fibers in the (+) or (-) buckled regime for up to 30 min (Methods).

As with buckled naked DNA, we found that topo II was extremely processive also on the buckled chromatin substrates. Fig. 5b shows all traces of the constant-extension measurements on chromatin fibers, in both the (+) and (-) buckled states. As before, most traces did not show any permanent loss of activity, although a few traces contained long pauses (> 30 s duration). We again conservatively classified individual traces as being “active” until only the first such pause, and fitting the active fractions over time to exponentials yielded lower bounds on the mean activity lifetimes of 112 ± 29 min in the (+) buckled state and 169 ± 96 min in the (-) buckled state (Fig. 5c). Similarly, we obtained lower bounds on the processivities of 12400 ± 4700 turns in the (+) buckled state and 22400 ± 7600 turns in the (-) buckled state (Fig. 5d). Comparison with the processivity for buckled naked DNA (Fig. 2d) suggests that chromatinization enhances topo II processivity under high torsional stress.

The activity lifetimes and processivities are similar for (-) and (+) buckled states of chromatin. However, we maintained the (-) buckled state at approximately -30 turns of supercoiling and the (+) buckled state at approximately +60 turns of supercoiling state, due to the highly asymmetric extension-supercoiling state relation of chromatin. Thus, the chromatin was maintained under topological states with similar levels of compaction and torsion but different magnitudes of supercoiling. If we were to maintain the chromatin at +30 turns to match the magnitude of supercoiling of the (-) buckled state, then the chromatin would be within the extension plateau regime (Fig. 5a).

We then examined the effect of chromatinization on topo II activity under low torsional stress, using the repeated winding experiment (Fig. 2b) but with the same chromatin fibers (Fig. 6; Methods). For this measurement, we again pre-bound topo II to chromatin fibers that satisfied our selection criteria (Supplementary Fig. 8; Methods) and repeatedly checked for topo II activity by adding or removing a small number of turns (Methods). As before, if topo II was active on a chromatin fiber after a rewinding step, the extension of the tether increased as the added supercoils were relaxed. Fig. 6a shows a typical trace of the repeated winding experiment with chromatin in the (+) winding direction, in which +20 turns were added every 5 minutes and topo II remained active for the full 30 minutes of the measurement. Most chromatin fibers were stable throughout this experiment, with only 1 nucleosome lost on average in 30 min (Supplementary Fig. 9).

**Figure 6.**
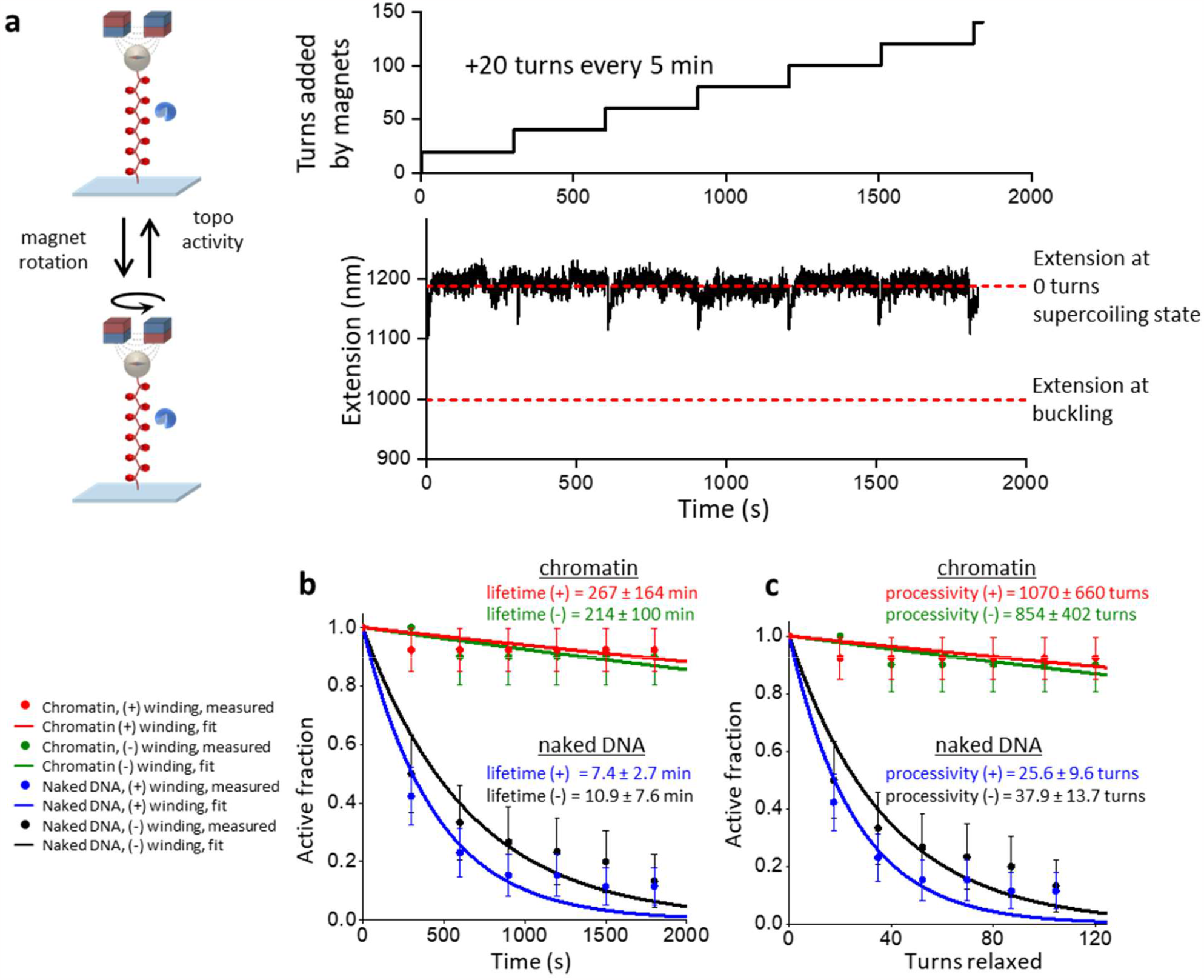
Chromatinization enhances topo II activity under low torsional stress. (**a**) Measurements of topo II activity lifetime and processivity on chromatin under low torsional stress. 20 turns were added every 5 min to chromatin fibers by rotation of the magnets. After each winding step, chromatin fibers with active topo II were identified by increases in extension. In the example trace, the chromatin fiber was under (+) winding, and its extension was consistently greater than that at the buckling (the lower dashed red line) for the entire active time duration, indicating that topo II relaxation occurred within the pre-buckled regime. (**b**) Activity lifetime of topo II on chromatin under low (+) torsional stress (*N* = 13) and low (-) torsional stress (*N* = 10) and 0.5 pN magnetic force; error bars represent counting error. These data were fit to exponential curves to estimate the activity lifetimes. For comparison, we also show the corresponding data and fits for the same DNA substrates containing 64 repeats of a nucleosome-positioning element but without nucleosomes under low (+) torsional stress (*N* = 28) and low (-) torsional stress (*N* = 30). (**c**) Processivity of topo II on chromatin under low (+) and (-) torsional stress from the same set of traces considered in b; error bars represent counting error. These data were fit to exponential curves to estimate the processivities.

As shown in Figs. 6b and 6c, even under low torsional stress, chromatin supported long activity lifetimes (267 ± 164 min for (+) winding; 214 ± 100 min for (-) winding) and processivities (1070 ± 660 turns for (+) winding; 854 ± 402 turns for (-) winding), both of which are about 20-fold greater than those of their naked DNA counterparts (compare Fig. 6 and Fig. 2). These results demonstrate that chromatin stabilizes topo II activity under low levels of torsional stress. Unlike a torsionally relaxed naked DNA with fewer DNA crossings, a torsionally relaxed chromatin fiber naturally contains DNA crossings because of the juxtaposition of the entry and exit DNA segments at each nucleosome^40^, which may give rise to the stabilization of topo II activity on chromatin under low torsional stress. In addition, this property predicts that increased torsional stress on chromatin should further increase the juxtapositions and facilitate topo II activity, which may contribute to the even greater activity lifetimes and processivities we observed on buckled chromatin.

### The relaxation of chromatin by topo II is influenced by compaction

The existence of three distinct topological states in the chromatin extension-supercoiling state relationship suggests that the relaxation rate of topo II may depend on the topological state of a chromatin substrate; however, this dependence has not yet been measured directly. Single-molecule methods used in previous studies are not suitable for such measurements because of the nonlinearity in the buckled regimes combined with the shallowness of the slope in the extension plateau regime^27^. To directly measure topo II relaxation rate in these three regimes of chromatin, we used the constant-extension method with chromatin fibers containing ∼50 nucleosomes (Fig. 7a; Methods). For each trace, a chromatin fiber that satisfied our selection criteria (Supplementary Fig. 8; Methods) was bound with a single molecule of topo II and held for 2 min each in: 1) the (-) buckled regime at 50% of its maximum extension, 2) the midpoint of the extension plateau regime, and 3) the (+) buckled regime at 50% of its maximum extension. As before, the mean rate of topo II relaxation in each state was calculated from the turns added by the magnets during the corresponding clamping step.

**Figure 7.**
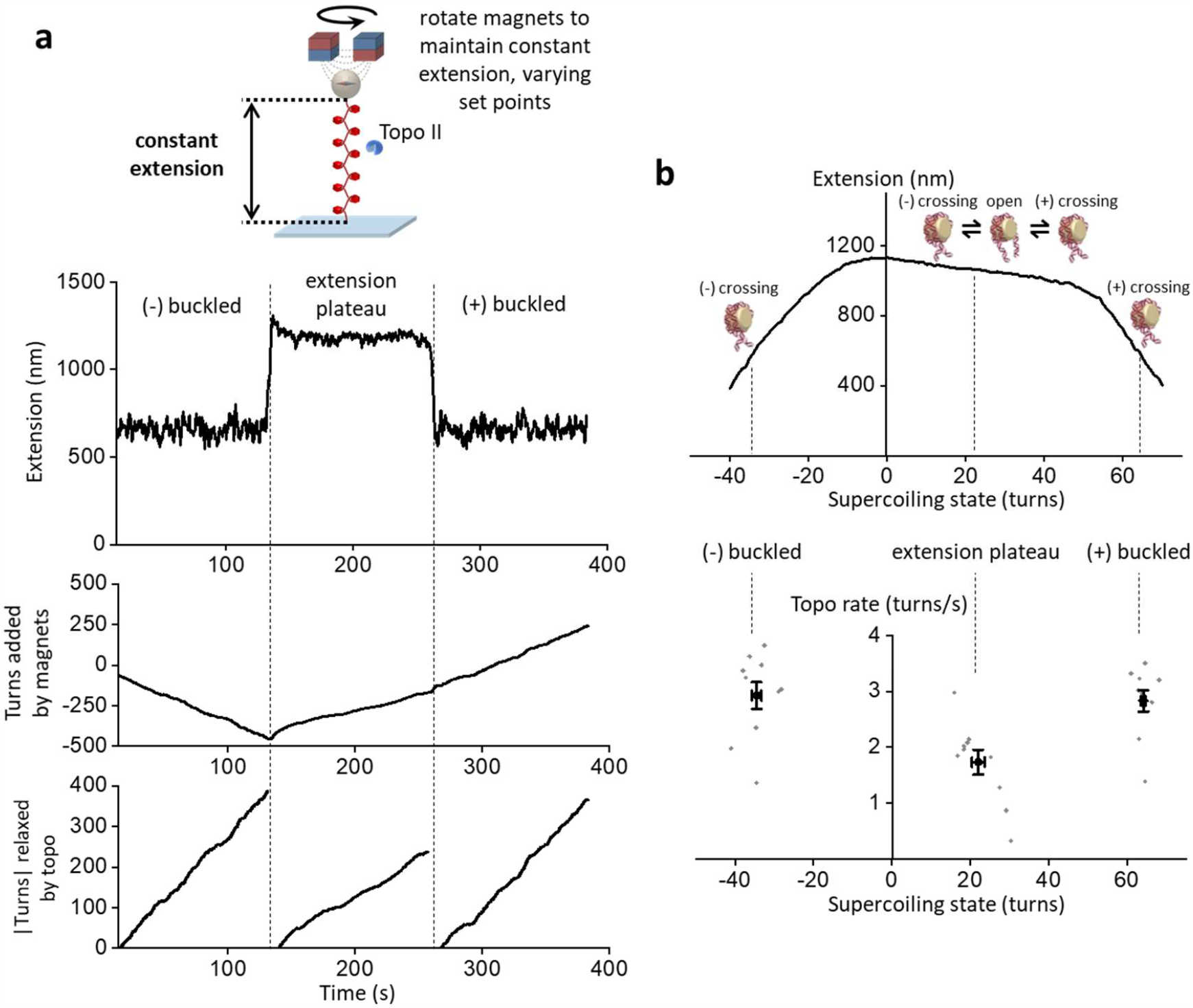
Topo II relaxation of chromatin depends on compaction. (**a**) Measurements of topo II relaxation rate on chromatin fibers in different topological states. A chromatin fiber with an active topo II was held for 2 min in each of the following states: the (-) buckled state at 50% of its maximal extension, then at the midpoint of the extension plateau regime, then finally in the (+) buckled state at 50% of its maximal extension. The mean rate of topo II in each topological state was calculated from the number of turns relaxed by the bound topo II in the step corresponding to that topological state. (b) Topo II rate on chromatin fibers as a function of supercoiling state (*N* = 10). Both individual data points (gray) as well as their means and SEMs for each topological state (black) are shown. For clarity, the chromatin extension-supercoiling state relation is also shown.

We found that the magnitudes of the topo II relaxation rates in the (+) and (-) buckled regimes were not significantly different from each other but that the rate was slower in the extension plateau regime than in the buckled regimes by ∼40% (Fig. 7b). This result suggests that DNA crossings were less available for topo II activity on chromatin in the less compact extension plateau regime. On buckled chromatin, the mean rate was consistent with that on buckled naked DNA (Supplementary Figs. 1 and 3), indicating that topo II relaxation was not limited by the availability of DNA crossings in either of the buckled regimes.

Interestingly, our results suggest that both the shape of the chromatin extension-supercoiling state relationship and the rate of topo II activity on chromatin are dictated by the left-handed solenoidal DNA wrapping that is characteristic of nucleosomes^40^. Due to this chirality, the entry and exit DNA segments of an individual nucleosome can cross in a supercoiling-dependent manner, with three distinct states: a (+) or (-) state, in which the entry and exit DNA segments cross with the corresponding chirality, and an open state, in which the entry and exit DNA segments do not cross^38, 41, 42, 43^. In the (+) and (-) buckled regimes of the chromatin extension-supercoiling state relationship, individual nucleosomes reside primarily in the corresponding crossed state. By comparison, chromatin in the extension plateau regime comprises a mixture of nucleosomes in the (+), (-), or open states, with the partitioning determined by the degree of supercoiling. The extension plateau is a result of the gradual conversion of nucleosomes from the (-) state to the open and (+) states to absorb (+) supercoiling. Although the (+) and (-) states readily provide a crossing for topo II activity, the enzyme must rely on spontaneous looping to form a crossing for the open state. Since not all nucleosomes are in a crossed state in the extension plateau regime, topo II activity was relatively slower here as compared to the buckled regimes.

## DISCUSSION

Taken together, our results highlight how the dynamics of supercoiled DNA and chromatin substrates, into which new insights are still emerging^27, 28, 31, 39, 44, 45, 46^, impact topo II activity. Specifically, our data suggest that substrate dynamics dictate the processivity and rate of topo II activity by determining the availability of DNA crossings for recognition by topo II (Fig. 8).

**Figure 8.**
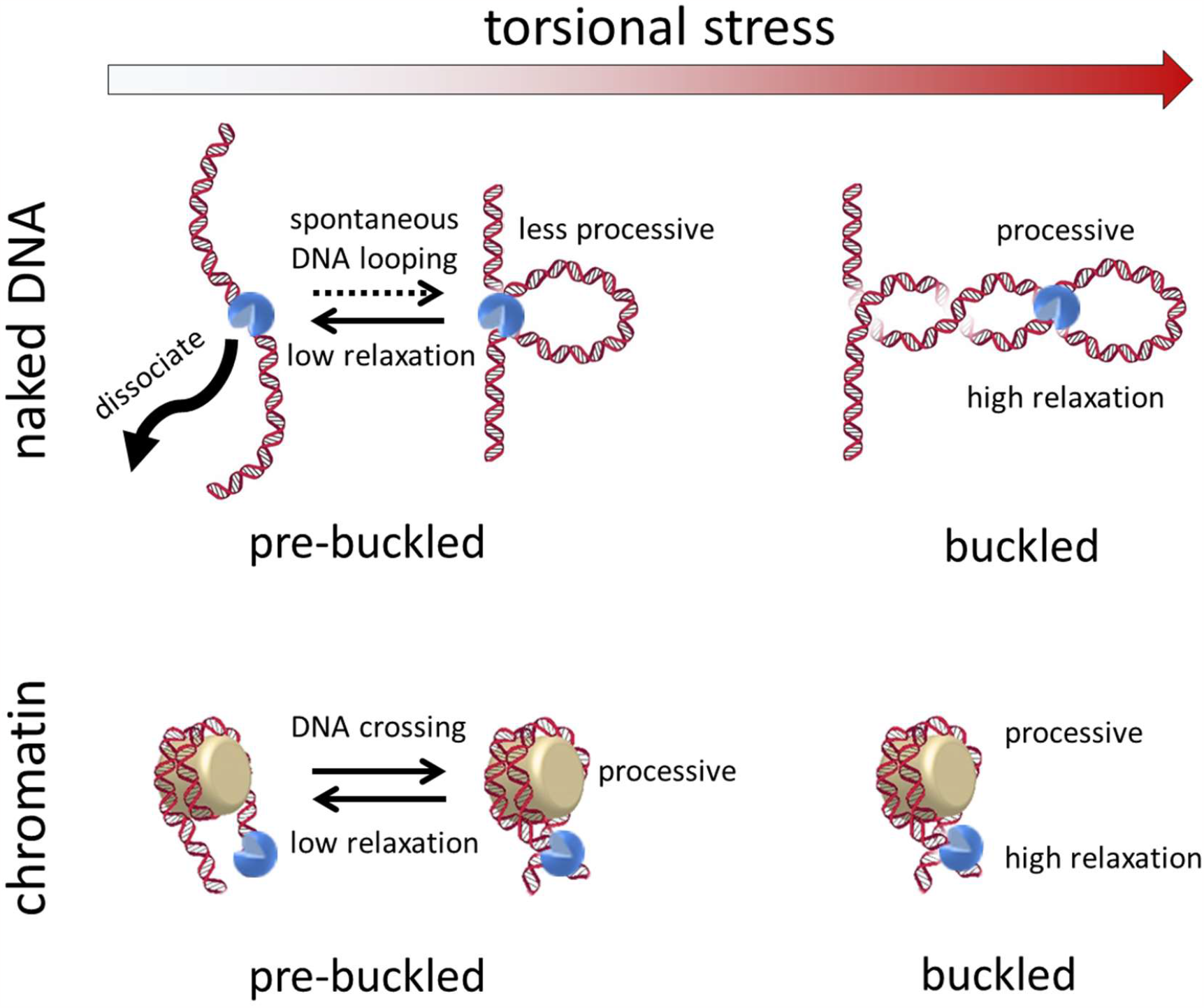
Chromatinization modulates topoisomerase II processivity. Topo II can capture spontaneous loops for activity on pre-buckled naked DNA under low torsional stress, but with greatly reduced activity lifetime, processivity, and rate compared to activity on buckled naked DNA under high torsional stress. Chromatinization enhances topo II activity lifetime and processivity for substrates under both low and high torsional stress, but the rate of activity is still reduced compared to that on chromatin under high torsional stress.

We unexpectedly found that topo II relaxed buckled DNA with much greater activity lifetime and processivity (Figs. 2 and 3) than previous estimates. Our data show that for naked DNA under high torsional stress, DNA crossings are readily available for persistent and efficient topo II activity. *In vivo*, buckled DNA is formed by high levels of torsional stress, which can arise from both transcription and replication. Our estimated lower bound of the processivity of a single molecule of topo II on buckled DNA far exceeds the necessary value to fully relax all of the supercoiling resulting from transcription of a typical gene (∼1.5 kb on average in yeast^47^) or replication between adjacent origins (∼30 kb on average in yeast^48, 49^). These processes result in ∼150 or ∼3000 turns of supercoiling on average, respectively, both of which are far less than the processivity measured by our experiments (> 6000 turns). Thus, a topo II that is actively relaxing supercoiling from either of these processes will almost certainly continue as long as the DNA substrate is under high torsional stress. During transcription, the high processivity of topo II on buckled DNA may enable topo II to counteract the increased transcription that occurs during gene activation and transcriptional bursting^50, 51^.

We also found that topo II could relax naked pre-buckled DNA, although with greatly reduced processivity compared to its activity on buckled DNA. Although previous single-molecule experiments were not sensitive to this capability, we were able to detect and characterize topo II activity in the pre-buckled regime (Fig. 4). We show that a simple model relating this activity to the probability of spontaneous loop formation can predict our measurements, suggesting that topo II captures (and may even help induce the formation of) spontaneous loops to relax pre-buckled DNA.

Although previous studies have examined topo II activity on nucleosomal substrates^27, 52, 53^, this work provides the first single-molecule examination of how topo II activity depends on the topological state of chromatin. Our data show that chromatinization greatly enhances topo activity lifetime and processivity (Figs. 5 and 6). The intrinsic DNA crossings afforded by nucleosomes within a torsionally relaxed chromatin fiber may contribute to the increased activity stability. The persistent activity of topo II with chromatin may enable topo II to maintain superhelical density at levels near zero as a housekeeping function throughout the genome and during steady-state transcription. In addition, the increased activity lifetime of topo II may ensure that the enzyme remains bound to daughter chromatin fibers under low torsional stress to promote decatenation after replication termination. This inherent stability raises the intriguing possibility that the dissociation of topo II from chromatin may need to be assisted by motor proteins, such as RNA polymerases, replisomes, or chromatin remodeling factors.

We also directly measured the rate of topo II relaxation on single chromatin fibers in different topological states (Fig. 7). Our results show that topo II relaxation rates are slower on chromatin under low torsional stress and faster on buckled, more compact chromatin substrates under high (-) or (+) torsional stress, consistent with topo II needing to capture DNA crossings for activity on chromatin. Taken together with the chromatin extension-supercoiling state relation, this rate variation suggests that the rate depends on the conformations of individual nucleosomes within the fibers. *In vivo*, the relatively slow relaxation of (+) supercoiling in pre-buckled chromatin downstream of an active RNA polymerase may permit supercoiling-induced destabilization of nucleosomal barriers ahead of the polymerase, while the faster relaxation of buckled chromatin by topo II may prevent transcription from stalling under high torsional stress^54^.

Collectively, our work elucidates the characteristics of topo II relaxation of both naked DNA and chromatin substrates and provides insights into how topo II responds to the complex torsional profile of DNA that likely occurs within a cell. Although this study primarily focused on *S. cerevisiae* topo II, vertebrates possess two topoisomerase II isoforms, IIα and IIβ, which are structurally and biochemically similar but still serve distinct roles *in vivo*^*1, 2, 3, 55*^. As all type IIA topoisomerases share the same core mechanism, the properties of yeast topo II revealed in this study may be consistent in all type IIA topoisomerases, including those in higher organisms; consistent with this idea, we found that human topoisomerases IIα and IIβ also show exceptionally long activity lifetimes and processivities on buckled DNA (Fig. 3). Future studies complementing the methods described in this work with fluorescence techniques could broaden the utilities of these single-molecule assays and further demonstrate the general importance of substrate dynamics in controlling fundamental biological processes.

## METHODS

### DNA template construction

The 12.7 kb DNA construct is composed of a 12,688 bp center segment flanked by ∼500 bp multi-labeled adapters at each end, as previously described^28^. In brief, the center segment was PCR amplified from λ DNA using primers designed with 2-4 mismatched nucleotides to engineer restriction enzyme recognition sites. The resulting DNA was double digested and spin column purified to produce a center segment with overhangs for ligation.

The adapters for the 12.7 kb DNA construct were created as previously described^27^. In brief, a 482 bp DNA sequence was created from low nucleosome affinity sequences identified by a large-scale nucleosome occupancy study^56^. This sequence was then cloned into pUC57 to form pNFRTC, and the adapters were then PCR amplified from this plasmid with 22% of dATP replaced by biotin-14-dATP (Invitrogen 19524016) for the biotin adapter and 22% of dTTP replaced by digoxigenin-11-dUTP (MilliporeSigma 11093088910) for the digoxigenin adapter. The labeled PCR products were spin column purified, restriction digested, and ligated to the ends of the center segment. The sample was gel purified to remove unligated adapters.

The 6.5 kb DNA construct is composed of a 6,481 bp center segment flanked by ∼500 bp multi-labeled adapters at each end. First, λ DNA was digested with BamHI-HF (NEB R3136) and KpnI-HF (NEB R3142), after which the largest fragment (11,552 bp) was gel purified. This fragment was then inserted into pUC19 (NEB N3041) to form pL14225 (a.k.a. pMDW133). A 6,546 bp sequence was then PCR amplified from this plasmid using LongAmp Taq DNA Polymerase and one primer designed with one mismatched nucleotide to engineer a restriction enzyme recognition site (primer sequences 5’-GCCACCTGACGTCTAAGAAACCATTATTATCA-3’ and 5’-GC<u>A</u>GGTCCTTTCCGGCAATCAGG-3’, mismatch is underlined). The PCR product was spin column purified and then digested with BssSI-v2 and PpuMI (NEB R0560) in order to form the 6,481 bp center segment with 4 or 3 bp overhangs on the ends, respectively.

The ∼500 bp multi-labeled adapters for the 6.5 kb DNA construct were created similarly to those for the 12.7 kb DNA construct. The adapters were PCR amplified from pNFRTC with Taq DNA polymerase and primers designed with 2 or 3 mismatches (forward primers 5’-GCTTCA<u>C</u>TCGT<u>G</u>CTTTTGTTCCTTATTTT-3’ for the biotin adapter and 5’-CGTTGTAAAA<u>G</u>GAC<u>CC</u>CCAGTGAAT-3’ for the digoxigenin adapter, mismatches are underlined; reverse primers were the same as those for the adapters in the 12.7 kb construct) with the same fraction of labeled dNTPs as the adapters for the 12.7 kb DNA construct. The PCR products were spin column purified and digested with BssSI-v2 or PpuMI before ligation to the 6.5 kb center segment. The sample was then gel purified before use.

The torsionally constrained 64-mer DNA construct was formed as previously described^27^. In brief, the construct is composed of a 12,667 bp center segment containing 64 tandem repeats of a sequence containing the 601 nucleosome positioning element (NPE)^57^ and ∼500 bp multi-labeled adapters at each end. The center segment was formed by inserting the 64 tandem repeats into pUC19^58^, double digesting, and then spin column purifying. The ∼500 bp multi-labeled adapters for the 64-mer construct were formed as previously described^27^. In brief, the adapters were PCR amplified with 25% of dATP replaced by biotin-14-dATP or 25% of dTTP replaced by digoxigenin-11-dUTP. The labeled PCR products were restriction digested and ligated to the center segment of the 64-mer DNA construct.

### Protein purification

Yeast topoisomerase II, human topoisomerase IIα, and human topoisomerase IIβ were purified as previously described^59^. In brief, tagged topoisomerases were expressed in *S. cerevisiae*, which were harvested by centrifugation, flash frozen dropwise in liquid nitrogen, and lysed by cryogenic grinding. The tagged protein was purified from the lysate by a series of purification selection columns with the tags removed in an intermediate step. The purified proteins were filter-concentrated and flash-frozen for storage.

Histone octamers were purified as previously described^27, 60, 61, 62, 63^. In brief, nuclei were extracted from HeLa-S3 cell pellets (National Cell Culture Center HA.48) in Nuclear Pellet Prep Lysis Buffer^64^. Core histones were purified using a hydroxyapatite Bio-gel HTP gel slurry (Bio-Rad Laboratories) as previously described^65^ but with the omission of MNase digestion before fractionation. The purified histones were stored at -80 °C.

### Torque measurements using the angular optical trap

The torque measurements were carried out with our in-house created angular optical trap (AOT)^24, 25^. The AOT permits simultaneous measurement and control of force, extension, torque, and rotation of DNA via nanofabricated quartz cylinders^25, 27, 28^. A cylinder may be rotated via rotation of the trapping laser polarization, and the torque on the cylinder is directly measured via a change in the angular momentum of the laser after interaction with the cylinder. For the data shown in Fig. 1, the extension was filtered by a 0.05 s sliding window, and the torque was filtered by a 2 s sliding window. These experiments were performed in the topo reaction buffer (10 mM Tris-HCl pH 8.0, 50 mM NaCl, 50 mM KCl, 3 mM MgCl_2_, 0.1 mM EDTA, 1 mM DTT, 0.5 mM TCEP, 1 mM ATP, and 1.5 mg/ml β-casein).

### Sample chamber preparation

Sample chambers were prepared as previously described^27^ with the listed modifications.

In brief, microscope coverslips were coated with a thin layer of nitrocellulose by spin coating. Coated coverslips were rinsed and stored in a clean plastic container for 48 h or more before being assembled into a microfluidic sample chamber using inert silicone high vacuum grease (Dow Corning 1597418). In contrast to previous work, nitrocellulose-coated coverslips were used for both the top and bottom surfaces of the sample chamber instead of only the bottom surface.

Tethers for MT experiments were formed as previously described^27^ with some modifications. First, a nitrocellulose coated sample chamber was incubated with 10 ng/μL anti-digoxygenin (Vector Labs MB-7000) in phosphate buffered saline (PBS) for 30 min. Next, 1 μm magnetic beads (Dynabeads MyOne Streptavidin T1, ThermoFisher 65601) coated with biotin and digoxigenin labeled DNA were bound to the surface to serve as fiducial markers. Then, 5 mg/ml β-casein (MilliporeSigma C6905) was incubated in the chamber for at least 1 h to passivate the surfaces. The DNA substrate of interest was then incubated at concentrations ranging from 1.5 pM to 10 pM for 15 min, followed by magnetic beads at a concentration of 20 μg/ml for 15 min. The buffer in the sample chamber was then exchanged for the topo reaction buffer.

### Magnetic tweezers

All experiments were carried out on a custom-built magnetic tweezers (MT) setup based on previous designs^27, 66, 67, 68^. In brief, a pair of 0.25” cube neodymium dipole magnets (K&J Magnetics B444) with a separation gap of 0.5 mm maintained by aluminum spacers was used to generate the magnetic field over the stage of a microscope. Magnetic bead images were collected via a 20x/0.75 NA air objective lens (Nikon Plan Apo 20x 0.75 NA) onto a 2.3 MP camera (Basler acA01920-155um) at a frame rate of 40 fps and an exposure time of 0.75 ms. Magnetic bead positions were tracked in three dimensions with LabVIEW as previously described^27, 69^.

### Selection of a DNA tether with a single active topo II

Immediately prior to each topoisomerase experiment, 0.5 pM of topo II in topo reaction buffer was introduced into the chamber and incubated for 2 min at room temperature. The sample chamber was then flushed with 5x chamber volume of the topo reaction buffer without topo II, then flushed again after 5 min. These two washes reduce further topo II binding events during measurements. After this process, typically ∼10-15% of our tethers showed topo II relaxation activity when wound down, which we defined as “active.” Because of this low probability of a tether being initially active, the initial activity from these tethers was most likely from a single topo II molecule. However, during the course of the subsequent measurement, an additional topo II molecule, which could be a consequence of incomplete flushing or topo II unbound from other tethers, could bind to these tethers and relax them. To determine the likelihood of this occurrence, we performed the conventional “wind down and wait” control experiment^19^ after topo II binding and flushing and monitored the tethers that were initially inactive but became active later in the measurements. We found this probability to be ∼ 17 ± 5% during the entire 30 min experiment. For a shorter duration of 30 s (the pause threshold used to define the activity duration), this probability was only ∼ 0.3 ± 0.1%. Therefore, our measurements should be predominately from a single topo II. Furthermore, we found that the active tether fraction decreased from 70% to 10-15% with the two additional washing steps, indicating that the washes are effective at removing free topo II. In addition, experiments with the constant-extension method provided a natural method to select out traces with multiple topo II molecules, so we were able to impose more stringent selection criteria to ensure a single topo II enzyme condition. In this method, the onset of activity from an additional topo II molecule caused an abrupt doubling in the relaxation rate. Thus, we could readily identify and exclude those traces from further analysis. All experiments were performed at 23°C.

For experiments that required the selection of one naked DNA tether with an active topo II, tethers were initially wound down by adding +60 turns by magnet rotation at 10 turns/s. 10 s after winding, a tether was chosen randomly from the tethers with extension increase greater than 250 nm. For experiments that required the selection of one chromatin fiber with an active topo II, tether selection was performed similarly but with the tethers initially wound down by adding -30 turns instead. In this way, we limited possible structural deformation of nucleosomes that could be induced by (+) torsion^30^.

### Implementation of the constant-extension clamp

Supplementary Fig. 1 explains the overall operation of the constant-extension clamp. We show a schematic flow chart of the extension feedback control and algorithm (Supplementary Fig. 1b). The feedback control was implemented as a proportional controller in LabVIEW. Given a setpoint extension, the error between the setpoint and the extension of a selected tether was calculated for each frame. Every five frames, the mean of the error was calculated for the previous five frames then multiplied by the gain of the proportional controller (0.05 turns/s/nm for all experiments) in order to set the magnet rotation rate. We also show representative raw traces of how the clamp responds to bead position changes (Supplementary Fig. 1c, d) and validation of the method by the agreement of the topo II relaxation rate of post-buckled DNA measured with the constant-extension method and the conventional method (Supplementary Fig. 1e).

### Calculation of pre-buckled supercoiling state reached by topo II

In the repeated winding experiments to probe topo II activity on pre-buckled DNA (Figs. 2 and 4; Supplementary Fig. 2), the DNA was buckled immediately after each winding step. The measured extension at this time point was used to determine DNA supercoiling state in the preceding step (as in Supplementary Fig. 2a). To do so, we used the mean extension over 1 s after rewinding and the extension-supercoiling state relation to determine the supercoiling state after the rewinding step, subtracted it from the number of turns added in the rewinding step, and added a small number of turns to account for topo II activity on buckled DNA during the 0.875 s rewinding step and 1 s averaging after rewinding at the measured rate (described below in the next section). The resulting value provides the supercoiling-state of the DNA in the proceeding step.

### Calculation of topo II rate on buckled DNA from repeated winding experiment

In the repeated winding experiment used to measure topo II activity on pre-buckled DNA (Fig. 4), the rate of topo II activity on buckled DNA was calculated while the DNA was still buckled after each winding step (before relaxation to the buckling transition). The extension data in this regime was first filtered by a 5 s sliding window and converted to supercoiling state by using the initially measured extension-supercoiling state relation. The rate was then calculated for each relaxation step from linear fits to the supercoiling state data. The mean of such rates calculated from each force condition was used for the corresponding linear extrapolation of buckled topo II rates on pre-buckled DNA (Fig. 4, dashed blue lines).

### Direct conversion of extension to supercoiling state of pre-buckled DNA

For the direct conversion of the measured extension to supercoiling state in the repeated winding experiment used to characterize relaxation of pre-buckled DNA by topo II (Fig. 4b), a mean extension versus time curve was obtained by averaging all relaxation traces after alignment at the buckling transition. The supercoiling state versus time curve was then determined from this extension versus time curve using the extension-supercoiling state relation at this force.

### Coarse-Grained Monte Carlo Simulations of DNA

The DNA was modeled as a discrete worm-like chain. Equilibrium configurations of naked DNA under external force and torsion were then generated using Monte-Carlo (MC) simulations with a Metropolis-Hastings procedure^70^ as described previously^71^, with parameters shown in Table S1. As the model does not directly incorporate the twisting of individual chain segments, the overall twist change ∆*Tw* in the DNA was indirectly computed by computing the global writhe change ∆*Wr* of the chain^72^ and invoking the conservation of linking number ∆*Lk* = ∆*Tw* + ∆*Wr*.

Using a scheme described in a recent study^73^, we implemented two types of structural moves to effectively sample a wide range of DNA configurations as part of the Metropolis process: local chain moves (pivot^74^ and crankshaft^75^ rotations) and full chain exchanges between neighboring replicas^76^. For the local moves, we sampled the degree of pivot and crankshaft rotations from a uniform distribution between [-50°,50°]. To implement the full chain exchanges, 12 parallel simulations were simultaneously generated on separate CPU cores. MC samples were then generated by applying 1000 local move trials followed by replica-exchange trials between configurations with neighboring linking numbers, ∆*Lk*_i_ and ∆*Lk*_i+1_, whose modified Metropolis criterion is given by the differential energy term:

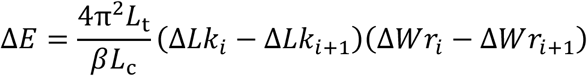

where 1/*β* is the thermal energy, *L*_t_ = 109 nm is the twist persistence length, and *L*_c_ is the contour length of the DNA (Table S1). The configurations were saved after five successful replica-exchange trials between any pair of neighboring simulations, and the process was repeated until there were 25,000 saved independent configurations for each simulation.

Since MC simulations allow for segments to cross one another, such moves could lead to creation of knots. Hence, a knot-checking algorithm^77, 78^ was implemented after every 1000 moves, and any conformation that contained a knot was rejected. To facilitate equilibration of long DNA molecules, we parallelized the writhe computation algorithm^72^ and used MATLAB Parallel Computing Toolbox to increase the overall sampling speed. To confirm that thermal equilibrium was achieved, we plotted the simulated DNA extension-supercoiling state relations and compared them to the experimentally measured extension-supercoiling state relations. Once the equilibriums were attained, subsequent saved DNA configurations were analyzed to estimate the average rate of DNA crossing formation.

We used a previously described method^73^ to obtain the kinetics of DNA crossing formation from our saved DNA conformations. First, we computed the distribution of all site-to-site distance measurements *R* between every single pair of non-contiguous vertices of all our saved conformations. Next, the distributions were transformed into effective free energy profiles that were used to model one-dimensional diffusion along the distance reaction coordinate *R* and estimate the average time it takes to reach from an “uncrossed” state to a state with a crossing. In our analysis, a crossing (sometimes called a DNA juxtaposition^79^) is defined as a state where two segments of the same DNA molecule are within a maximal threshold *R*_T_ = 10 nm distance of each other. Finally, the mean first passage time ⟨*T*⟩ for crossing formation for a given superhelical density σ and force *F* can be estimated using:

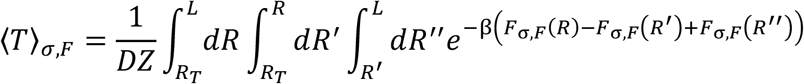

where 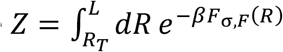 is the uncrossed state partition function, *F* _*σ,F*_ (*R*) is the effective free energy profile along the reaction coordinate *R* for a given set of *σ* and *F*, and *D* is the effective diffusion coefficient. We then used the average rate of DNA crossing 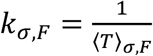 (Supplementary Fig. 6) to model the supercoiling relaxation activity of topo II by using a Michaelis-Menten type equation to model the rate of supercoiling relaxation by topo II:

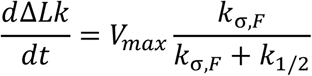

where *V*_max_ and *k*_1/2_ are constants that could be obtained by comparing to experimental data. Specifically, *V*_max_ was obtained experimentally from the post-buckling relaxation rate in the repeated winding experiment (3.4, 3.1, and 3.0 turns/s for 0.5, 1.1, and 1.6 pN, respectively) and *k*_1/2_ = 6.8 ± 1.3 loops/s was a global fitting parameter computed by integrating the theoretical relaxation rate over time and minimizing the chi-squared difference between the integrated theoretical curves and experimental data (Fig. 4b).

To examine how DNA bending by topo II may affect the DNA crossing rate, we repeated the simulations at *F* = 0.5 pN with the introduction of a rigid bend in the center of the DNA (Supplementary Fig. 7a). The geometry of this bend structure is based on a two-kink model^35, 80^, where the DNA is kinked in two positions, creating a flat-bottomed ‘V’ with a total bend angle of 120 degrees. The average looping rate was calculated in the same way as before (Supplementary Fig. 7b), and the results were fitted to the same Michaelis-Menten type equation. We found that the simulations including a DNA bend could be consistent with our results, but with *k*_1/2_ increased to 14.6 ± 3.9 loops/s. The greater value of *k*_1/2_ reflects the increased formation of DNA loops at low values of superhelical density, allowing for more efficient loop capture by topo II to match our experimental results (Supplementary Fig. 7c).

### Nucleosome assembly

Nucleosome arrays were assembled onto the 64-mer DNA construct and assayed as previously described^27^. In brief, histones were deposited onto the 64-mer DNA by gradient salt dialysis from 2.0 M to 0.6 M over 18 h at 4 °C at different molar ratios (0.25:1 to 2.5:1) of histone octamers to 601 DNA repeats^61, 81^. An equal mass of 147 bp random-sequence competitor DNA was included to avoid nucleosome over-assembly^27^. The quality and saturation level of the final product were assayed by gel electrophoresis in 0.7% native agarose and then by stretching with an optical tweezers setup.

### Chromatin fiber characterization on MT

Prior to each topoisomerase experiment on a chromatin substrate, we performed a twisting experiment as previously described^27^ to measure the extension-supercoiling state relation of the substrate under a constant force of 0.5 pN in the range -40 to +70 turns (Figs 5a and 7b). These extension-supercoiling state relations were fit to five-piece functions composed of three linear regions joined by two parabolic regimes (Supplementary Fig. 8a) ^27^. From these fits, we obtained the peak extension and the turns of the left and right “buckling-like” transitions 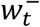 and 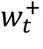, which were defined to be the intercepts of the middle linear regime with the left and right linear regimes, respectively^27^. We compared these parameters with those of chromatin fibers that were well-saturated with primarily full nucleosomes (as opposed to subnucleosomal structures, such as tetrasomes), which we previously characterized with the nucleosome disruption assay described below. This enabled selection of chromatin fibers with acceptable saturation and composition using only the initial extension-supercoiling state relation, without disrupting the nucleosomes.

To benchmark the nucleosome saturation and composition of a chromatin substrate, we performed a separate nucleosome disruption assay. After measuring initial extension-supercoiling state relations as described above, we disrupted the nucleosomes in the fiber with a combination of high stretching force and high ionic strength and interpreted the resulting extensions based on a previously-established multi-stage disruption model of chromatin^27, 61, 62, 82, 83, 84^. Specifically, we examined the extension of each chromatin fiber at 2 pN, at which the “outer-turn” DNA (interacting primarily with the H2A/H2B dimers) remains bound to the histone core octamers, and 6 pN, at which the outer-turn DNA has been mostly released. The outer-turn DNA is ∼72 bp in length for each nucleosome, so the number of nucleosomes initially containing a wrapped outer-turn *N*_out_ could be determined from the DNA length released. Next, we fully dissociated nucleosomes with a 5 min incubation in 2 M NaCl followed by an additional wash with the same salt concentration. This process released the “inner-turn” DNA (interacting primarily with the H3/H4 tetramer) from the histone core octamers, which is ∼75 bp in length for each nucleosome. The number of bound tetramers *N*_in_ could be determined from the total DNA released by the high salt wash, and the length of the underlying naked DNA template could be measured after all nucleosomes were fully dissociated. These experiments were performed using nucleosome array samples assembled with varying molar ratios of histone octamers to 601 DNA repeats to obtain data over a broad range of saturations.

To establish benchmark relationships between nucleosome saturation/composition and the shape of the extension-supercoiling state relation using the data from the nucleosome disruption assay, we first considered the relative values of *N*_out_ and *N*_in_. We only considered fibers satisfying (|*N*_in_ − *N*_out_|/*N*_in_) ≤ 0.15, which indicates *N*_out_ and *N*_in_ being comparable within measurement uncertainties, as having an acceptable composition of nucleosomes. For these fibers, we estimated the total number of full nucleosomes *N*_nuc_ = (*N*_out_ + *N*_in_)/2 and plotted the relationship between the peak extensions from their initial extension-supercoiling state relations and *N*_nuc_ (Supplementary Fig. 8b, data points). A linear fit to this relationship (Supplementary Fig. 8b, dashed black line) was later used to calculate *N*_nuc_ for each fiber prior to experiments with topo II. In addition, we plotted the relationship between the positions of the buckling-like transitions 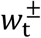 and *N*_nuc_ for nucleosome fibers with acceptable composition (Supplementary Fig. 8c, data points). Linear fits to the relationships between 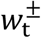 and *N*_nuc_ (Supplementary Fig. 8c, dashed black lines) were later used to evaluate the composition of nucleosomes in each fiber prior to experiments with topo II.

For experiments with topo II, we only selected fibers with maximal extension consistent with 50 ± 5 nucleosomes and with 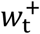 and 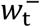 within the 95% confidence intervals of the linear fits to 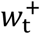 and 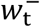 versus *N*_nuc_ (Supplementary Fig. 8c, green regions). These selection criteria removed tethers with extreme nucleosome saturation, poor nucleosome composition, short underlying DNA (i.e. loss of repeats in the underlying 64-mer DNA), or partial sticking of the fiber to a surface. For the fibers that passed selection with ∼50 nucleosomes, 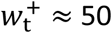 turns, suggesting that each nucleosome absorbs about 1 turn before buckling.

## ACKNOWLEDGEMENTS

We thank members of the Wang Laboratory and Drs. Z.J. Wang and G. Lambert for helpful discussion and comments. We especially thank Drs T.T. Le and X. Gao for technical advice and generation of the naked DNA templates used in this study and Dr. J.P. Sethna for technical advice regarding the Monte-Carlo simulations in this study. This work is supported by the National Institutes of Health grants R01GM136894 (to M.D.W.), T32GM008267 (to M.D.W.), and R01-CA077373 and R35-CA263778 (to J.M.B.). M.D.W. is a Howard Hughes Medical Institute investigator.

## AUTHOR CONTRIBUTIONS

J.L., J.T.I., and M.D.W. designed single-molecule assays. J.L., M.W., J.T.I., and Y.H. performed experiments. G.S. performed simulations. J.L., M.W., and Y.H. analyzed data. J.T.I. constructed the MT with contribution from J.L. on software. J.H.L. J.J., and J.M.B. purified topo II and characterized it biochemically. R.M.F. purified histones. S.P. and M.W. prepared and characterized the chromatinized DNA substrates. J.L., G.S., and M.D.W. wrote the initial draft, and all authors contributed to manuscript revisions. M.D.W. supervised the project.

## References

1. Champoux JJ. DNA topoisomerases: structure, function, and mechanism. Annual review of biochemistry 70, 369–413 (2001).

2. Pommier Y, Sun Y, Shar-yin NH, Nitiss JL. Roles of eukaryotic topoisomerases in transcription, replication and genomic stability. Nature reviews Molecular cell biology 17, 703–721 (2016).

3. Vos SM, Tretter EM, Schmidt BH, Berger JM. All tangled up: how cells direct, manage and exploit topoisomerase function. Nature reviews Molecular cell biology 12, 827–841 (2011).

4. Wang JC. Cellular roles of DNA topoisomerases: a molecular perspective. Nature reviews Molecular cell biology 3, 430–440 (2002).

5. Berger JM, Gamblin SJ, Harrison SC, Wang JC. Structure and mechanism of DNA topoisomerase II. Nature 379, 225–232 (1996).

6. Roca J, Wang JC. DNA transport by a type II DNA topoisomerase: evidence in favor of a two-gate mechanism. Cell 77, 609–616 (1994).

7. Lee JH, Berger JM. Cell cycle-dependent control and roles of DNA topoisomerase II. Genes 10, 859 (2019).

8. Kouzine F, Sanford S, Elisha-Feil Z, Levens D. The functional response of upstream DNA to dynamic supercoiling in vivo. Nat Struct Mol Biol 15, 146–154 (2008).

9. Kouzine F, et al. Transcription-dependent dynamic supercoiling is a short-range genomic force. Nat Struct Mol Biol 20, 396–403 (2013).

10. Keszthelyi A, Minchell NE, Baxter J. The Causes and Consequences of Topological Stress during DNA Replication. Genes (Basel) 7, (2016).

11. Goto T, Wang JC. Cloning of yeast TOP1, the gene encoding DNA topoisomerase I, and construction of mutants defective in both DNA topoisomerase I and DNA topoisomerase II. Proceedings of the National Academy of Sciences 82, 7178–7182 (1985).

12. Brill SJ, DiNardo S, Voelkel-Meiman K, Sternglanz R. Need for DNA topoisomerase activity as a swivel for DNA replication for transcription of ribosomal RNA. Nature 326, 414–416 (1987).

13. Kim RA, Wang JC. Function of DNA topoisomerases as replication swivels in Saccharomyces cerevisiae. Journal of molecular biology 208, 257–267 (1989).

14. Baxter J, Diffley JF. Topoisomerase II inactivation prevents the completion of DNA replication in budding yeast. Molecular cell 30, 790–802 (2008).

15. Charvin G, Bensimon D, Croquette V. Single-molecule study of DNA unlinking by eukaryotic and prokaryotic type-II topoisomerases. Proceedings of the National Academy of Sciences 100, 9820–9825 (2003).

16. McClendon AK, et al. Bimodal recognition of DNA geometry by human topoisomerase IIα: preferential relaxation of positively supercoiled DNA requires elements in the C-terminal domain. Biochemistry 47, 13169–13178 (2008).

17. Roca J, Wang JC. The probabilities of supercoil removal and decatenation by yeast DNA topoisomerase II. Genes to Cells 1, 17–27 (1996).

18. Seol Y, Gentry AC, Osheroff N, Neuman KC. Chiral Discrimination and Writhe-dependent Relaxation Mechanism of Human Topoisomerase IIα. Journal of Biological Chemistry 288, 13695–13703 (2013).

19. Strick TR, Croquette V, Bensimon D. Single-molecule analysis of DNA uncoiling by a type II topoisomerase. Nature 404, 901–904 (2000).

20. McClendon AK, Rodriguez AC, Osheroff N. Human topoisomerase IIα rapidly relaxes positively supercoiled DNA: implications for enzyme action ahead of replication forks. Journal of Biological Chemistry 280, 39337–39345 (2005).

21. Osheroff N, Shelton ER, Brutlag DL. DNA topoisomerase II from Drosophila melanogaster. Relaxation of supercoiled DNA. J Biol Chem 258, 9536–9543 (1983).

22. Zechiedrich EL, Osheroff N. Eukaryotic topoisomerases recognize nucleic acid topology by preferentially interacting with DNA crossovers. The EMBO journal 9, 4555–4562 (1990).

23. Roca J, Berger JM, Wang JC. On the simultaneous binding of eukaryotic DNA topoisomerase II to a pair of double-stranded DNA helices. J Biol Chem 268, 14250–14255 (1993).

24. La Porta A, Wang MD. Optical torque wrench: angular trapping, rotation, and torque detection of quartz microparticles. Phys Rev Lett 92, 190801 (2004).

25. Deufel C, Forth S, Simmons CR, Dejgosha S, Wang MD. Nanofabricated quartz cylinders for angular trapping: DNA supercoiling torque detection. Nat Methods 4, 223–225 (2007).

26. Forth S, Deufel C, Sheinin MY, Daniels B, Sethna JP, Wang MD. Abrupt buckling transition observed during the plectoneme formation of individual DNA molecules. Phys Rev Lett 100, 148301 (2008).

27. Le TT, et al. Synergistic coordination of chromatin torsional mechanics and topoisomerase activity. Cell 179, 619-631. e615 (2019).

28. Gao X, Hong Y, Ye F, Inman JT, Wang MD. Torsional stiffness of extended and plectonemic DNA. Physical review letters 127, 028101 (2021).

29. Baird CL, Harkins TT, Morris SK, Lindsley JE. Topoisomerase II drives DNA transport by hydrolyzing one ATP. Proc Natl Acad Sci U S A 96, 13685–13690 (1999).

30. Bancaud A, et al. Nucleosome chiral transition under positive torsional stress in single chromatin fibers. Molecular cell 27, 135–147 (2007).

31. Marko JF, Neukirch S. Competition between curls and plectonemes near the buckling transition of stretched supercoiled DNA. Physical Review E 85, 011908 (2012).

32. Sankararaman S, Marko JF. Formation of loops in DNA under tension. Physical Review E 71, 021911 (2005).

33. Moroz JD, Nelson P. Torsional directed walks, entropic elasticity, and DNA twist stiffness. Proc Natl Acad Sci U S A 94, 14418–14422 (1997).

34. Dong KC, Berger JM. Structural basis for gate-DNA recognition and bending by type IIA topoisomerases. Nature 450, 1201–1205 (2007).

35. Lee I, Dong KC, Berger JM. The role of DNA bending in type IIA topoisomerase function. Nucleic Acids Res 41, 5444–5456 (2013).

36. Lane AB, Gimenez-Abian JF, Clarke DJ. A novel chromatin tether domain controls topoisomerase IIalpha dynamics and mitotic chromosome formation. J Cell Biol 203, 471–486 (2013).

37. Sundararajan S, et al. Methylated histones on mitotic chromosomes promote topoisomerase IIalpha function for high fidelity chromosome segregation. iScience 26, 106743 (2023).

38. Bancaud A, et al. Structural plasticity of single chromatin fibers revealed by torsional manipulation. Nat Struct Mol Biol 13, 444–450 (2006).

39. Kaczmarczyk A, Meng H, Ordu O, Noort Jv, Dekker NH. Chromatin fibers stabilize nucleosomes under torsional stress. Nature communications 11, 1–12 (2020).

40. Luger K, Mäder AW, Richmond RK, Sargent DF, Richmond TJ. Crystal structure of the nucleosome core particle at 2.8 Å resolution. Nature 389, 251–260 (1997).

41. De Lucia F, Alilat M, Sivolob A, Prunell A. Nucleosome dynamics. III. Histone tail-dependent fluctuation of nucleosomes between open and closed DNA conformations. Implications for chromatin dynamics and the linking number paradox. A relaxation study of mononucleosomes on DNA minicircles. Journal of molecular biology 285, 1101–1119 (1999).

42. Prunell A, Sivolob A. Paradox lost: nucleosome structure and dynamics by the DNA minicircle approach. New Comprehensive Biochemistry 39, 45–73 (2004).

43. Sivolob A, Prunell A. Nucleosome conformational flexibility and implications for chromatin dynamics. Philosophical Transactions of the Royal Society of London Series A: Mathematical, Physical and Engineering Sciences 362, 1519–1547 (2004).

44. Desai PR, Brahmachari S, Marko JF, Das S, Neuman KC. Coarse-grained modelling of DNA plectoneme pinning in the presence of base-pair mismatches. Nucleic acids research 48, 10713–10725 (2020).

45. Dittmore A, Brahmachari S, Takagi Y, Marko JF, Neuman KC. Supercoiling DNA locates mismatches. Physical review letters 119, 147801 (2017).

46. van Loenhout MT, De Grunt M, Dekker C. Dynamics of DNA supercoils. Science 338, 94–97 (2012).

47. Dujon B. The yeast genome project: what did we learn? Trends in Genetics 12, 263–270 (1996).

48. Bell SP, Labib K. Chromosome duplication in Saccharomyces cerevisiae. Genetics 203, 1027–1067 (2016).

49. Sekedat MD, Fenyö D, Rogers RS, Tackett AJ, Aitchison JD, Chait BT. GINS motion reveals replication fork progression is remarkably uniform throughout the yeast genome. Molecular systems biology 6, 353 (2010).

50. Suter DM, Molina N, Gatfield D, Schneider K, Schibler U, Naef F. Mammalian genes are transcribed with widely different bursting kinetics. Science 332, 472–474 (2011).

51. Zenklusen D, Larson DR, Singer RH. Single-RNA counting reveals alternative modes of gene expression in yeast. Nat Struct Mol Biol 15, 1263–1271 (2008).

52. Fernández X, Díaz-Ingelmo O, Martínez-García B, Roca J. Chromatin regulates DNA torsional energy via topoisomerase II-mediated relaxation of positive supercoils. The EMBO journal 33, 1492–1501 (2014).

53. Salceda J, Fernández X, Roca J. Topoisomerase II, not topoisomerase I, is the proficient relaxase of nucleosomal DNA. The EMBO journal 25, 2575–2583 (2006).

54. Ma J, Wang M. Interplay between DNA supercoiling and transcription elongation. Transcription 5, e28636 (2014).

55. Linka RM, Porter AC, Volkov A, Mielke C, Boege F, Christensen MO. C-terminal regions of topoisomerase II α and II β determine isoform-specific functioning of the enzymes in vivo. Nucleic acids research 35, 3810–3822 (2007).

56. Kaplan N, et al. The DNA-encoded nucleosome organization of a eukaryotic genome. Nature 458, 362–366 (2009).

57. Lowary P, Widom J. New DNA sequence rules for high affinity binding to histone octamer and sequence-directed nucleosome positioning. Journal of molecular biology 276, 19–42 (1998).

58. Wu C, Read C, McGeehan J, Crane-Robinson C. The construction of customized nucleosomal arrays. Analytical biochemistry 496, 71–75 (2016).

59. Lee JH, Wendorff TJ, Berger JM. Resveratrol: A novel type of topoisomerase II inhibitor. Journal of Biological Chemistry 292, 21011–21022 (2017).

60. Brennan LD, Forties RA, Patel SS, Wang MD. DNA looping mediates nucleosome transfer. Nature communications 7, 1–8 (2016).

61. Brower-Toland B, Wacker DA, Fulbright RM, Lis JT, Kraus WL, Wang MD. Specific contributions of histone tails and their acetylation to the mechanical stability of nucleosomes. Journal of molecular biology 346, 135–146 (2005).

62. Brower-Toland B, Wang MD. Use of optical trapping techniques to study single-nucleosome dynamics. Methods in enzymology 376, 62–72 (2004).

63. Li M, et al. Dynamic regulation of transcription factors by nucleosome remodeling. Elife 4, e06249 (2015).

64. Schnitzler GR. Isolation of histones and nucleosome cores from mammalian cells. Current protocols in molecular biology 50, 21.25. 21-21.25. 12 (2001).

65. Wolffe AP, Ura K. Transcription of dinucleosomal templates. Methods 12, 10–19 (1997).

66. Lipfert J, Hao X, Dekker NH. Quantitative modeling and optimization of magnetic tweezers. Biophysical journal 96, 5040–5049 (2009).

67. Seol Y, Neuman KC. Magnetic tweezers for single-molecule manipulation. In: Single Molecule Analysis). Springer (2011).

68. Strick TR, Allemand J-F, Bensimon D, Bensimon A, Croquette V. The elasticity of a single supercoiled DNA molecule. Science 271, 1835–1837 (1996).

69. Lansdorp BM, Tabrizi SJ, Dittmore A, Saleh OA. A high-speed magnetic tweezer beyond 10,000 frames per second. Review of Scientific Instruments 84, 044301 (2013).

70. Metropolis N, Rosenbluth AW, Rosenbluth MN, Teller AH, Teller E. Equation of state calculations by fast computing machines. The journal of chemical physics 21, 1087–1092 (1953).

71. Vologodskii AV, Marko JF. Extension of torsionally stressed DNA by external force. Biophysical journal 73, 123–132 (1997).

72. Klenin K, Langowski J. Computation of writhe in modeling of supercoiled DNA. Biopolymers: Original Research on Biomolecules 54, 307–317 (2000).

73. Starr CH, Bryant Z, Spakowitz AJ. Coarse-grained modeling reveals the impact of supercoiling and loop length in DNA looping kinetics. Biophys J 121, 1949–1962 (2022).

74. Baumgärtner A, Binder K. Monte Carlo studies on the freely jointed polymer chain with excluded volume interaction. The Journal of Chemical Physics 71, 2541–2545 (1979).

75. Vologodskii AV, Levene SD, Klenin KV, Frank-Kamenetskii M, Cozzarelli NR. Conformational and thermodynamic properties of supercoiled DNA. Journal of molecular biology 227, 1224–1243 (1992).

76. Hukushima K, Nemoto K. Exchange Monte Carlo Method and Application to Spin Glass Simulations. Journal of the Physical Society of Japan 65, 1604–1608 (1996).

77. Frank-Kamenetskiĭ MD, Vologodskiĭ AV. Topological aspects of the physics of polymers: The theory and its biophysical applications. Soviet Physics Uspekhi 24, 679 (1981).

78. Virnau P. Detection and visualization of physical knots in macromolecules. Physics Procedia 6, 117–125 (2010).

79. Huang J, Schlick T, Vologodskii A. Dynamics of site juxtaposition in supercoiled DNA. Proc Natl Acad Sci U S A 98, 968–973 (2001).

80. Hardin AH, Sarkar SK, Seol Y, Liou GF, Osheroff N, Neuman KC. Direct measurement of DNA bending by type IIA topoisomerases: implications for non-equilibrium topology simplification. Nucleic Acids Res 39, 5729–5743 (2011).

81. Huynh VA, Robinson PJ, Rhodes D. A method for the in vitro reconstitution of a defined “30 nm” chromatin fibre containing stoichiometric amounts of the linker histone. Journal of molecular biology 345, 957–968 (2005).

82. Brower-Toland BD, Smith CL, Yeh RC, Lis JT, Peterson CL, Wang MD. Mechanical disruption of individual nucleosomes reveals a reversible multistage release of DNA. Proceedings of the National Academy of Sciences 99, 1960–1965 (2002).

83. Hall MA, Shundrovsky A, Bai L, Fulbright RM, Lis JT, Wang MD. High-resolution dynamic mapping of histone-DNA interactions in a nucleosome. Nature structural & molecular biology 16, 124–129 (2009).

84. Sheinin MY, Li M, Soltani M, Luger K, Wang MD. Torque modulates nucleosome stability and facilitates H2A/H2B dimer loss. Nature communications 4, 1–8 (2013).

