## Supplementary Information for "Chromatinization Modulates Topoisomerase II Processivity"

### Table of Contents:

**Supplementary Figures 1 – 9**

**Supplementary Table 1**

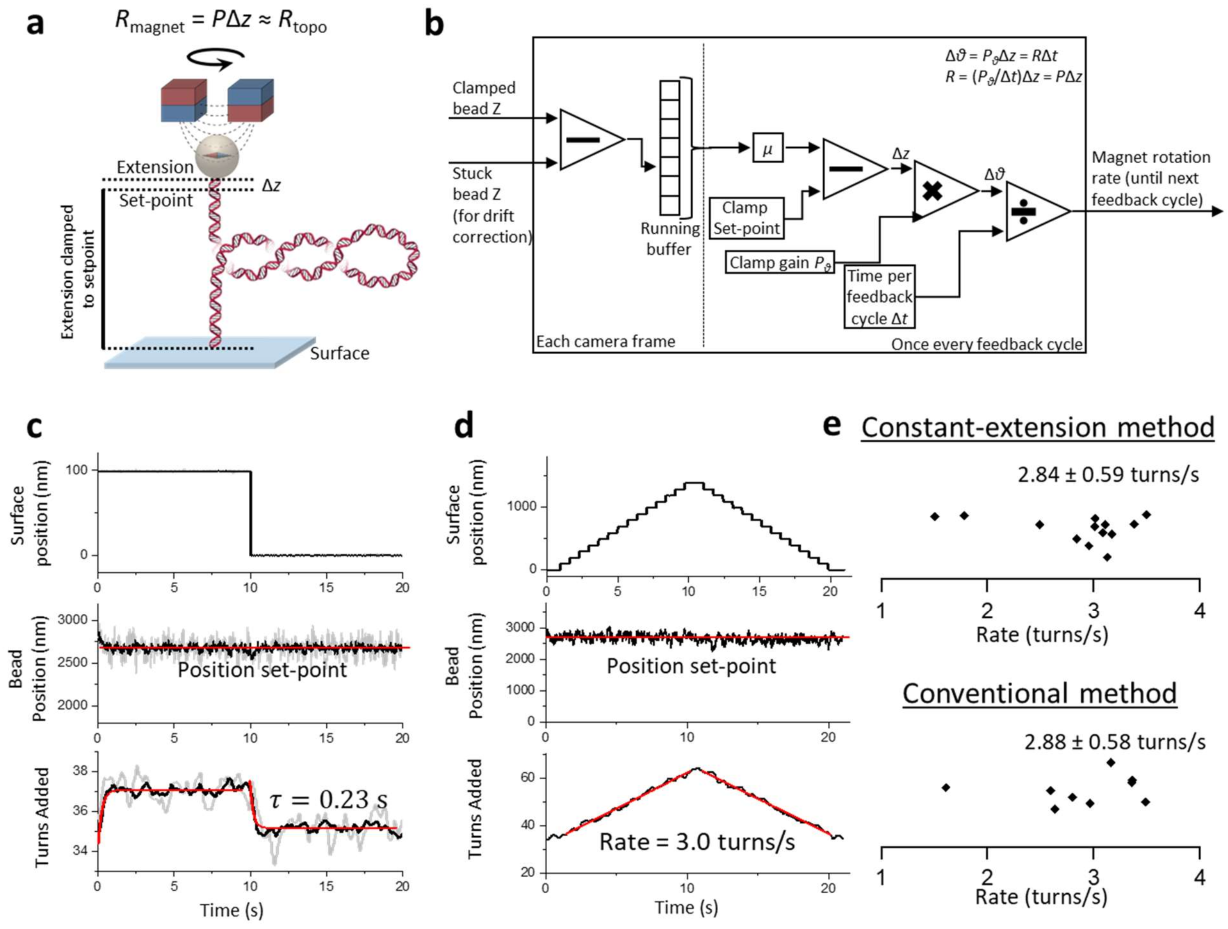


**Supplementary Figure 1. The constant-extension clamp.**

(**a**) Diagram of the constant-extension clamp applied to a DNA tether. The clamp is implemented as a proportional clamp with a gain $P$. The magnet rotation rate $R_{magnet}$ is calculated as a multiple of the difference extension signal $\Delta z$ between the measured extension of the DNA and the set point. When topo II relaxes the DNA with a rate $R_{\text{topo}}$, the clamp will rotate the magnets to keep the DNA extension close to the set point (close to $\Delta z = 0$) by matching the topo relaxation rate.

(**b**) Diagram of the magnet rotation rate calculation, using the measured $z$ positions of a tethered bead and a bead stuck to the coverslip surface (for drift correction) as input.

(**c**) Clamp response time. With the clamp off, each catalytic cycle of topo relaxation (2 turns) leads to a DNA extension change of ~100 nm for a buckled 12.7 kb DNA under 0.5 pN force. To determine the clamp response time, we mimicked a single catalytic cycle of topo II by stepping the surface with a piezo stage along $z$ in a 100 nm square-wave while clamping the bead's absolute position in the lab frame by magnet rotation and measured the bead position and turns added by the clamp over time (gray curve, single cycle); these data are also shown after averaging over 10 cycles from a single trace (black curve). The bead position remained close to the set-point while the magnet rotation reached an equilibrium with a response time $\tau$ = 0.23 ± 0.02 s (red, exponential fit).

(**d**) Measurement of the clamp response of a 12.7 kb buckled DNA held under 0.5 pN force to surface stepping of 100 nm every 0.67 s, mimicking supercoil relaxation by topo II at 3 turns/s. The bead's absolute position in the lab frame was clamped by magnet rotation. Data shown were from a single trace. The bead position stays near the position set-point for the entire measurement while the magnets rotated as 3.00 ± 0.02 turns/s, indicating that the clamp can measure the relaxation rate of topo II.

(**e**) Comparison of yeast topo II rates measured using the constant-extension clamp (*N* = 13, same traces as in Fig. 2) and the conventional method that uses post-buckling extension change (*N* = 9).

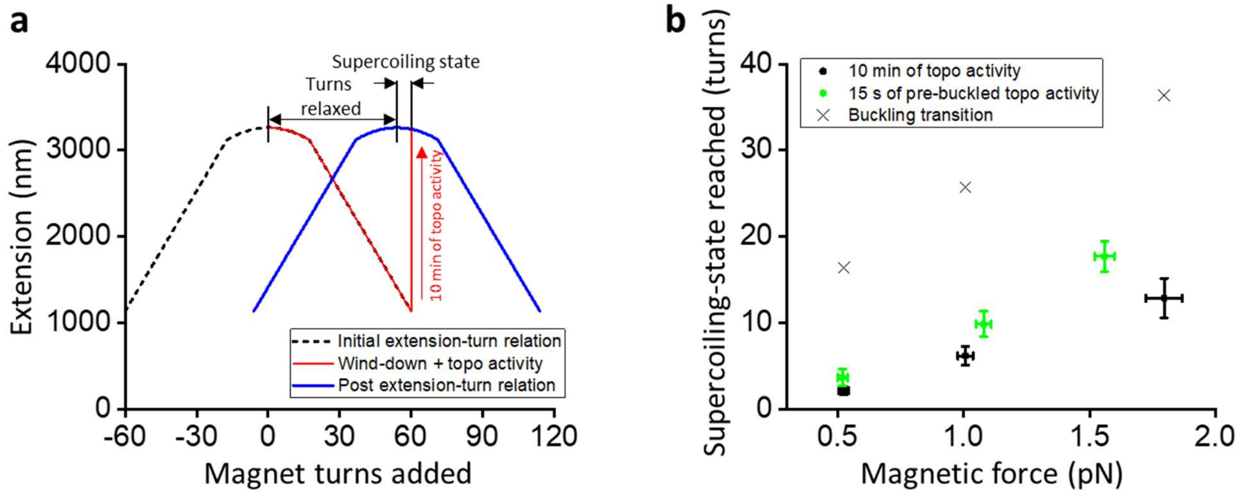


**Supplementary Figure 2. Topo II relaxation of pre-buckled DNA slows down sharply instead of stopping before reaching full relaxation.**

(**a**) Explanation on the determination of supercoiling state reached on pre-buckled DNA. After initial measurements of the extension-supercoiling state relation in the absence of topo II, +60 turns were applied by rotation of the magnets to the DNA tethers with active topo II and the extensions of the DNA were tracked for 10 min (red curve). At the end of the 10 min relaxation step, ATP was flushed out of the sample chamber and a post extension-supercoiling state relation was measured by rotation of the magnets from 0 to +120 turns (blue curve). The extension-supercoiling state relation of each DNA molecule relaxed by topo II was no longer centered at zero turns but instead at the number of turns relaxed because the activity of the enzyme changed the topological state of the DNA while the magnets were held fixed. The supercoiling states reached by topo II activity were indicated by the difference between +60 turns (turns initially applied by magnet rotation) and the center of the second extension-supercoiling state relation (number of turns relaxed by topo II activity).

(**b**) Mean supercoiling-states reached by topo II as a function of magnetic force after 10 min of topo II activity (black points, *N* = 16, 16, and 12 for 0.5 pN, 1.0 pN, and 1.8 pN respectively); error bars represent SEMs. The 10 min of topo II activity relaxed the molecules slightly further on average than the 15 s of activity that was allowed in the repeated winding experiment (green points; data from Fig. 4), indicating that topo II relaxation rate did not slow down to zero at 15 s.

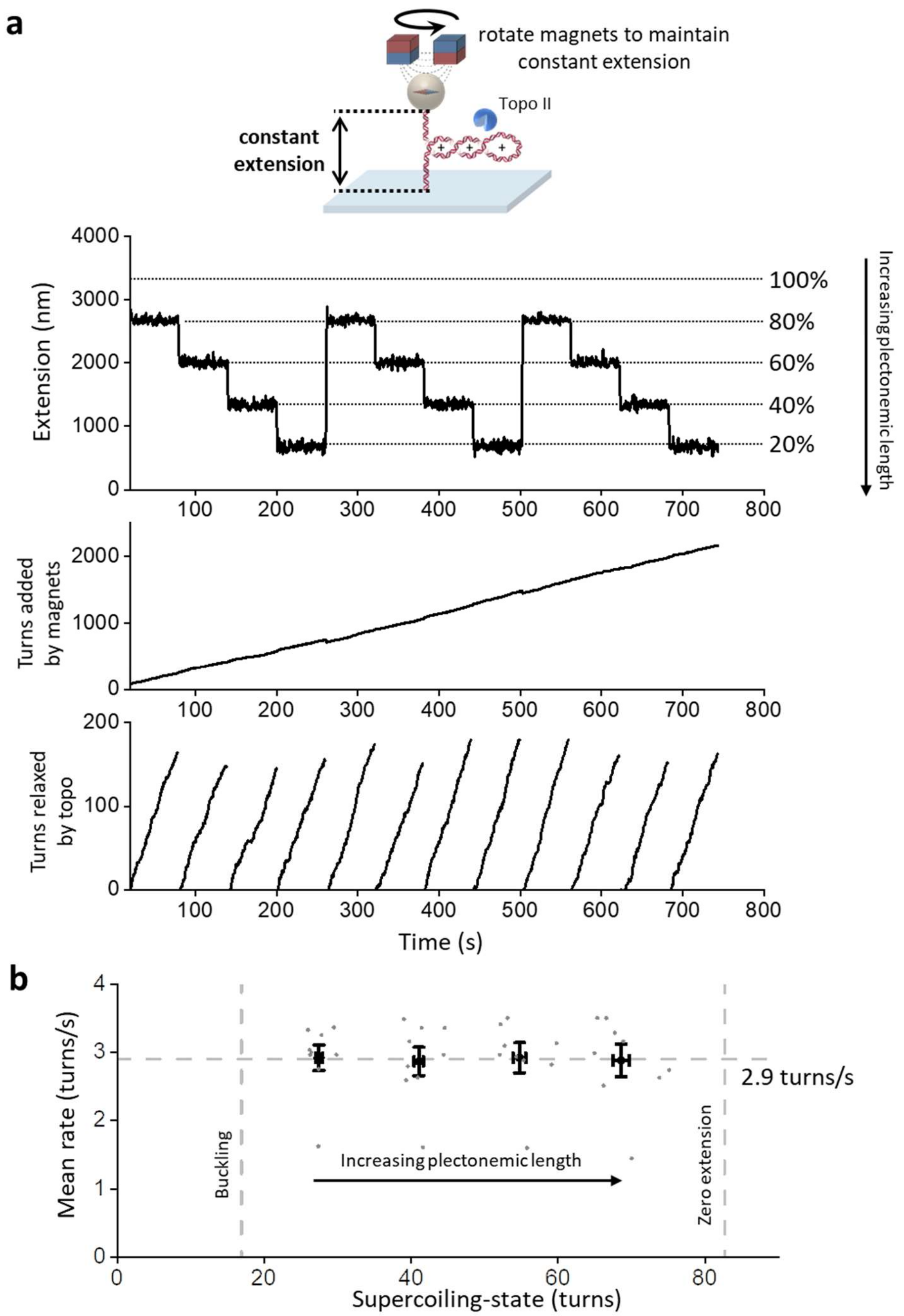


**Supplementary Figure 3**. **Buckling enhances topo II efficiency**.

(**a**) Measurements of topo II relaxation rate with varying plectonemic length. The extension of a 12.7 kb DNA molecule with an active topo II was clamped to 80%, 60%, 40%, and 20% of the maximal extension measured from the initial extension-supercoiling state relation for 60 s each using the constant-extension method. This sequence was repeated two more times for a total of three repetitions in each trace. The number of turns added by the magnets at each step was tracked over time, and the mean rate of topo II activity during each step was calculated from the number of turns relaxed by the bound topo II in that step.

(**b**) Mean topo II rate on buckled DNA as a function of supercoiling state under 0.5 pN force ($N$ = 9 traces). Individual data points are shown (gray) along with their means and SEMs for each cluster of supercoiling states (black). The horizontal line represents the mean value from all data points.

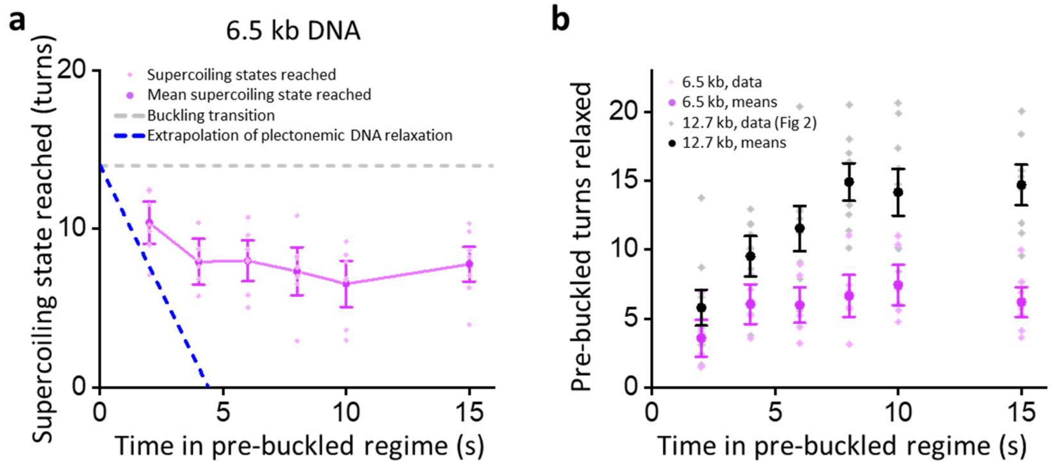


**Supplementary Figure 4. Topo II relaxation of pre-buckled DNA is slower for shorter DNA.**

(**a**) Supercoiling state reached by topo II relaxation into the pre-buckled regime as a function of the duration of topo II activity in the pre-buckled regime for 6.5 kb DNA under 1.1 pN magnetic force. Individual data points are shown as a scatter plot (light purple, $N = 7$ traces) along with their means and SEMs (dark purple). Purple lines connect the mean supercoiling states and are meant to guide the eye. This experiment was performed the same way as with the 12.7 kb DNA (Fig. 4) but with initial winding to +30 turns and additional winding steps of +20 turns each. The horizontal dashed gray line represents the supercoiling state of the buckling transition, and the dashed blue line is a linear extrapolation using the mean relaxation rate of buckled DNA.

(**b**) Mean number of pre-buckled turns relaxed as a function of the duration of topo II activity in the pre-buckled regime for 6.5 kb DNA (purple) and 12.7 kb DNA (black; from Fig. 4b) under 1.1 pN force; error bars represent SEMs; individual data points are shown in light purple for 6.5 kb DNA and gray for 12.7 kb DNA. The mean pre-buckled turns relaxed were calculated for each time point by taking the difference between the buckling transition and the mean supercoiling state reached.

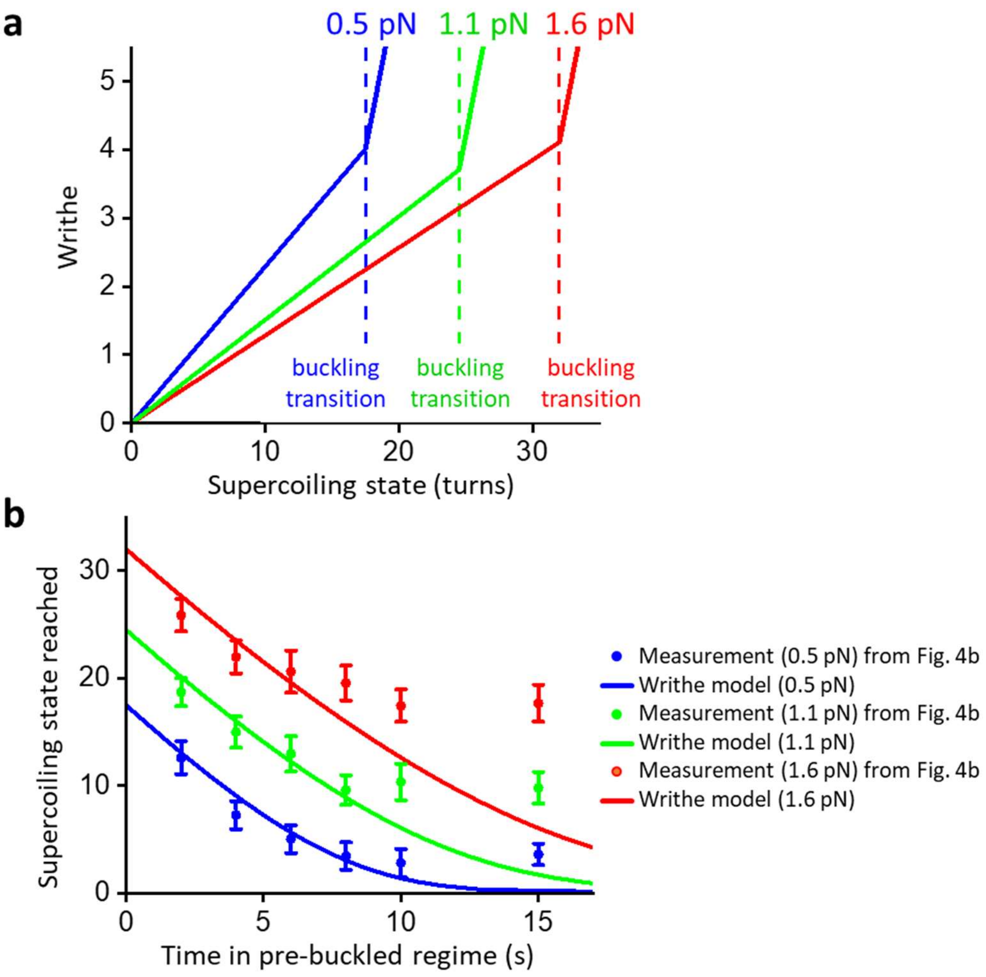


**Supplementary Figure 5. A simple writhe model cannot explain topo II relaxation activity on pre-buckled naked DNA.**

(**a**) In the pre-buckled regime, the linking number change $\Delta Lk$ partitions to twist change $\Delta Tw$ and writhe change $\Delta Wr$ according to the force-dependent effective torsional persistence length $C_{\text{eff}}(F)$ and the intrinsic torsional persistence length $C_0$: $\Delta Wr = [1 - C_{\text{eff}}(F)/C_0]\Delta Lk$. We have previously measured both $C_{\text{eff}}(F)$ and $C_0$ in the topo reaction buffer used in this work[1]. Using those values, we have obtained $\Delta Wr$ as a function of $\Delta Lk$ under three different forces for the 12.7 kb DNA used in Fig. 4.

(**b**) We have explored a simple writhe model to explain the relaxation of pre-buckled DNA by topo II, assuming that topo relaxation is only dependent on writhe according to the Michaelis-Menten relationship: $\frac{d\Delta Lk}{dt} = \frac{\Delta Wr}{\Delta Wr + \Delta Wr_0}$. Such a model has been previously used to explain the relaxation rate of topo II on post-buckled DNA[2]. This simple writhe model was used to fit the data from Fig. 4b. As shown in this figure, this model does not show a good agreement with the measurements obtained for the pre-buckled regime, suggesting that DNA writhing alone cannot explain the activity of topo II in this regime.

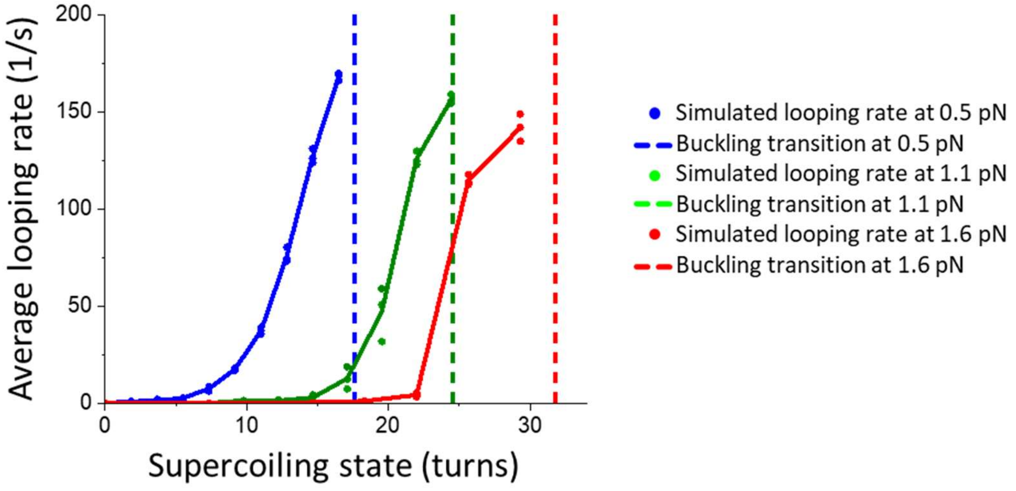


**Supplementary Figure 6. Simulated DNA looping rate as a function of turns added under varying forces.**

Simulated rate of DNA loop formation as a function of supercoiling state for 12.7 kb DNA under three stretching forces: 0.5 pN, 1.1 pN, and 1.6 pN. Using Monte Carlo simulation with the parameters shown in Supplementary Table 1 (Methods), equilibrium DNA configurations were generated for each force. The looping rate was obtained from three repeats of the simulation for each force (data points); lines connect the mean values of the three repeats shown to guide the eye.

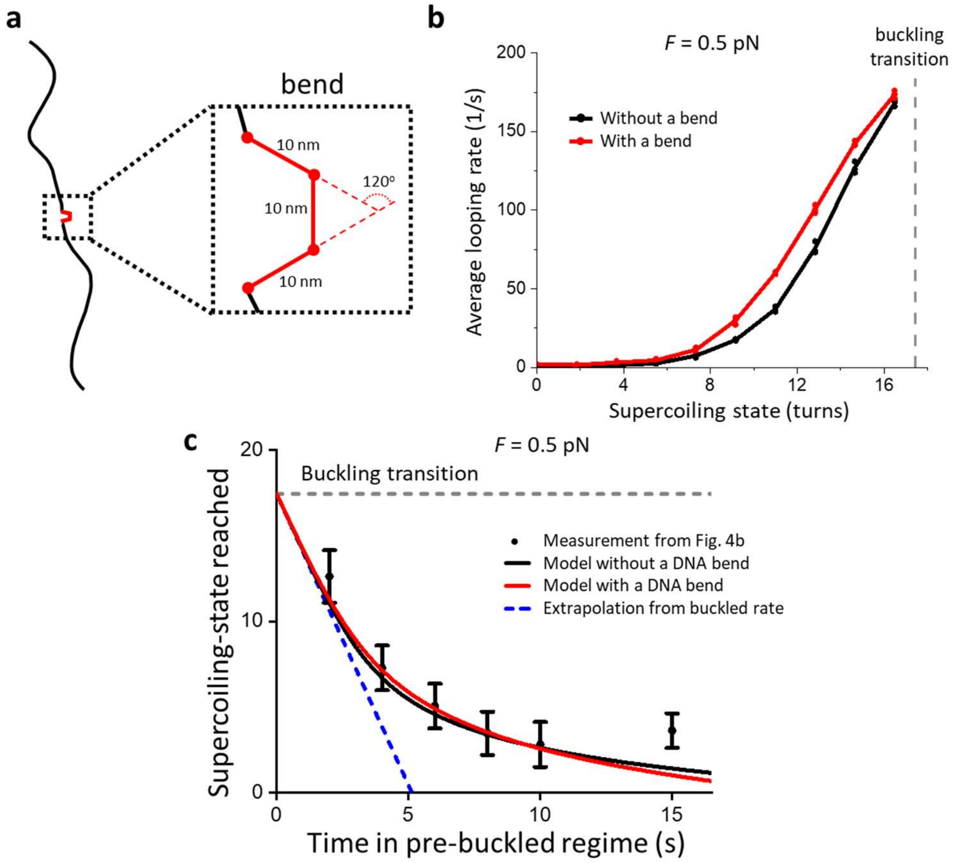


**Supplementary Figure 7. Monte Carlo simulation with consideration of a topo II-mediated DNA bend.**

(**a**) The geometry of a simulated DNA containing an intrinsic bend. The DNA bend is composed of two kinks to provide a total bend angle of 120 degrees to mimic the geometry of a topo II-mediated DNA bend.

(**b**) Simulated looping rate under 0.5 pN force on DNA with an intrinsic bend. The looping rate shown here was obtained from three repeats of the simulation (red data points), and the line connecting the mean values of the three repeats is shown to guide the eye. The simulation results without any intrinsic bend from Supplementary Fig. 6 are shown for comparison (black).

(**c**) Simulated pre-buckled relaxation of DNA under 0.5 pN force on DNA with an intrinsic bend (red curve). The simulation results without any intrinsic bend from Supplementary Fig. 6 are shown for comparison (black curve). Both simulations show good agreement with the measured topo relaxation activity in the pre-buckled regime (black data points).

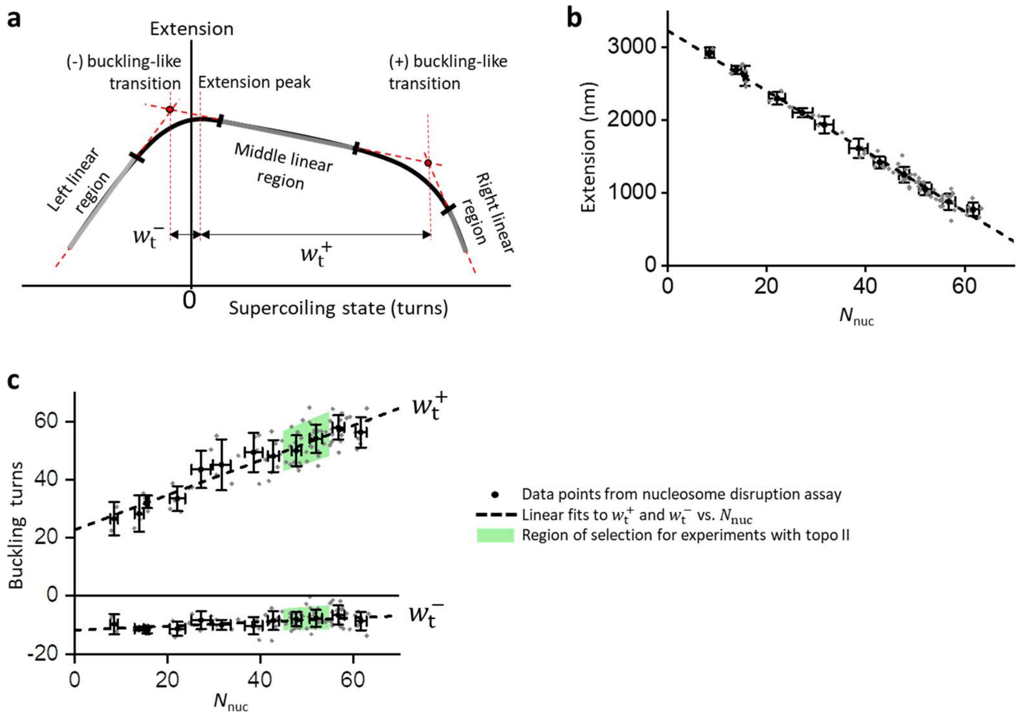


**Supplementary Figure 8. Chromatin fiber characterization and selection criteria for experiments with topo II.**

(**a**) 5-piece fit to the chromatin extension-supercoiling state relation (Methods). For each chromatin fiber, the extension at the peak of the extension-supercoiling state relation was used to calculate the number of nucleosomes $N_{\mathrm{nuc}}$. The positions of the (+) and (-) buckling-like transitions ($w_{\mathrm{t}}^{+}$ and $w_{\mathrm{t}}^{-}$, respectively) were used along with the peak extension to assess the quality of the chromatin fiber.

(**b**) Extension at the peak of the extension-supercoiling state relation as a function of $N_{\mathrm{nuc}}$ for chromatin fibers with acceptable composition (Methods). Individual data points are shown as gray data points. The data were binned by 5 nucleosome intervals with error bars representing

SDs (black data points). The linear fit (dashed line) was used to calculate $N_{\mathrm{nuc}}$ from the peak extension of each chromatin fiber used for experiments with topo II.

(**c**) $w_{\mathrm{t}}^{+}$ and $w_{\mathrm{t}}^{-}$ as functions of $N_{\mathrm{nuc}}$ for chromatin fibers with acceptable composition (Methods). Individual data points are shown as gray data points. The data were binned by 5-nucleosome intervals with error bars representing SDs (black data points). Linear fits (dashed lines) were used to evaluate the compositions of nucleosomes in each chromatin fiber. Chromatin fibers used for experiments with topo II (Figs. 5-7) were required to contain 50 ± 5 nucleosomes and have $w_{\mathrm{t}}^{\pm}$ within the 95% confidence intervals of linear fits to $w_{\mathrm{t}}^{\pm}$ versus $N_{\mathrm{nu}}$ (green regions).

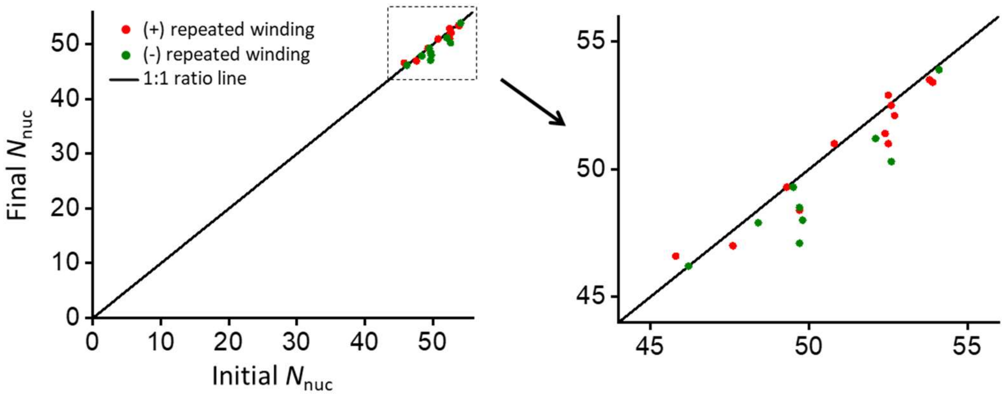


**Supplementary Figure 9. Chromatin fiber stability during the repeated winding experiments.**

Number of nucleosomes in chromatin fibers before (Initial $N_{nuc}$) and after (Final $N_{nuc}$) the repeated winding experiments shown in Fig. 6, calculated from extension-supercoiling state relations taken before and after the 30 min experiments for traces that did not show loss of topo II activity (traces that lost topo II activity were wound down and stuck to the surface). The right plot shows a zoomed in view of the region of interest. The mean nucleosome change was -0.4 ± 0.2 nucleosomes for (+) repeated winding ($N$ = 12 traces, mean ± SEM) and -1.1 ± 0.3 nucleosomes for (-) repeated winding ($N$ = 9 traces, mean ± SEM).

| Parameter | Value |
|---|---|
| DNA length | 12700 bp |
| Segment length | 10 nm |
| Temperature ($T$) | 293.0 K |
| Linear persistence length ($L_\mathrm{p}$) | 43 nm[3] |
| Twist persistence length ($L_\mathrm{t}$) | 109 nm[1] |
| Hard-wall cutoff for electrostatic repulsion | 5 nm[4] |

**Supplementary Table 1. Parameters used for MC simulation.**

These parameters were used with Monte Carlo simulation to generate equilibrium DNA configurations, from which the crossing probability was calculated as a function of supercoiling state for three different forces (Supplementary Figs. 6 and 7; Methods).

### References


1. Gao X, Hong Y, Ye F, Inman JT, Wang MD. Torsional stiffness of extended and plectonemic DNA. *Physical review letters* **127**, 028101 (2021).

2. Strick TR, Croquette V, Bensimon D. Single-molecule analysis of DNA uncoiling by a type II topoisomerase. *Nature* **404**, 901-904 (2000).

3. Wang MD, Yin H, Landick R, Gelles J, Block SM. Stretching DNA with optical tweezers. *Biophys J* **72**, 1335-1346 (1997).

4. Vologodskii AV, Marko JF. Extension of torsionally stressed DNA by external force. *Biophysical journal* **73**, 123-132 (1997).
